# SmdA is a novel cell morphology determinant in *Staphylococcus aureus*

**DOI:** 10.1101/2021.11.23.469651

**Authors:** Ine Storaker Myrbråten, Gro Anita Stamsås, Helena Chan, Danae Morales Angeles, Tiril Mathiesen Knutsen, Zhian Salehian, Volha Shapaval, Daniel Straume, Morten Kjos

## Abstract

Cell division and cell wall synthesis in staphylococci need to be precisely coordinated and controlled to allow the cell to multiply while maintaining their nearly spherical shape. The mechanisms ensuring correct placement of the division plane and synthesis of new cell wall have been studied intensively, however, hitherto unknown factors and proteins are likely to play key roles in this complex interplay. We here identified and investigated a protein with major influence on cell morphology in *Staphylococcus aureus*. The protein, named SmdA (for staphylococcal morphology determinant A), is a membrane-protein with septum-enriched localization. By CRISPRi knockdown and overexpression combined with different microscopy techniques, we demonstrate that proper levels of SmdA is necessary for cell division, including septum formation and cell splitting. We also identified conserved residues in SmdA that are critical for its functionality. Pulldown- and bacterial two-hybrid interaction experiments showed that SmdA interacts with several known cell division- and cell wall synthesis proteins, including penicillin binding proteins (PBPs) and EzrA. Notably, SmdA also affects susceptibility to cell wall targeting antibiotics, particularly in methicillin-resistant *S. aureus* (MRSA). Together, our results show that *S. aureus* is dependent on balanced amounts of membrane-attached SmdA in order to carry out proper cell division.

**Importance:** *Staphylococcus aureus* is an important human and animal pathogen. Antibiotic resistance is a major problem in treatment of staphylococcal infections, and cell division and cell wall synthesis factors have previously been shown to modulate susceptibility to antibiotics in this species. In the current work we investigated the function of an essential protein named SmdA, which was identified based on its septal localization and knockdown phenotype resulting in defective cellular morphologies. We demonstrate that this protein is critical for normal cell division in *S. aureus*. Depletion of SmdA sensitize resistant staphylococci to β-lactam antibiotics. This work thus reveals a new staphylococcal cell division factor and a potential future target for narrow spectrum antimicrobials or compounds to resensitize antibiotic resistant staphylococcal strains.

## Introduction

Most bacteria are surrounded by a shape-determining cell envelope which protects against lysis and interacts with the extracellular milieu. The cell envelope of the opportunistic, Gram-positive pathogen *Staphylococcus aureus* consists of a thick layer of peptidoglycan (PG) along with teichoic acids (TA) and cell wall-associated surface proteins. During a bacterial cell cycle, synthesis of PG and TA needs to be precisely regulated and coordinated with cell division, DNA replication and chromosome segregation. Tight control of these processes is critical for staphylococcal cells to maintain their integrity and nearly spherical shape as they multiply, and they are therefore attractive targets for antimicrobials (1). Exactly how such control is mediated in *S. aureus* is still not fully established, and hitherto unknown factors may be involved. In this work we describe a new staphylococcal cell morphology determinant.

Staphylococcal cell division is initiated by assembly of the Z-ring, consisting of polymerized FtsZ-proteins, that localizes to the future division septa (2). The Z-ring functions as a scaffold for cell division- and cell wall synthesis proteins which together constitute the divisome (3). Cell division in *S. aureus* occurs in alternating orthogonal planes, meaning that the new cell division plane is always perpendicular to the previous (4). Timely and spatial control of localization of the Z-ring assembly is most likely linked with chromosome segregation and DNA replication, involving proteins such as the nucleoid occlusion factor Noc, which ensures that the cells do not establish new septa across the chromosomes (5), and CcrZ, which connects initiation of DNA replication to cell division (6). The chromosomes and chromosome segregation also contribute to establish a physical barrier allowing the Z-ring only to be formed in an angle perpendicular to the previous division plane (4).

The Z-ring directs the synthesis of new PG in *S. aureus* to the septum. Synthesis of PG starts in the cytoplasm, where UDP-MurNAc-pentapeptide is first synthesized and then attached to the membrane by the enzyme MraY to form the PG precursor lipid I (7, 8). A GlcNAc residue and a pentaglycine side chain is attached to produce lipid II-Gly_5_ (9), which is flipped to the outer leaflet of the membrane by MurJ (10, 11) where it is incorporated into the existing PG mesh by transpeptidation (TP) and transglycosylation (TG) reactions. Specifically, the shape, elongation, division and sporulation (SEDS) proteins, FtsW and RodA with TG activity, work in pairs with monofunctional transpeptidases, the penicillin binding proteins PBP1 and PBP3, respectively (12, 13). While the PBP1-FtsW pair is essential and performs the septal cross wall synthesis, the non-essential PBP3-RodA pair is responsible for the slight elongation taking place in *S. aureus*. Additionally, *S. aureus* possesses two other PBPs; the bifunctional PBP2 with both TG and TP activity, whose role is essential in *S. aureus*, and the low-molecular weight PBP4, which controls the degree of PG crosslinks (14-16). Finally, MRSA strains have an additional PBP, PBP2A, a transpeptidase with low-affinity for β-lactam antibiotics (17, 18). In the final step of division, PG hydrolases break covalent bonds in PG for cell wall remodeling and daughter cell splitting. The major, bifunctional autolysin Atl, together with Sle1, for which expression are regulated by the two-component system WalKR, are the primary enzymes responsible for hydrolyzing the septal PG to allow splitting of daughter cells (19-22). The actual cross wall splitting is a mechanical process occurring within milliseconds (23, 24).

The spatiotemporal control of cell division and PG synthesis is directly and indirectly influenced by several factors. One of these is the anionic TA polymers, the second major component of the cell wall, which are either covalently linked to the PG (wall teichoic acids, WTA), or linked via a lipid-anchor to the plasma membrane (lipoteichoic acids, LTA). Mutations in enzymes involved in either WTA or LTA biosynthesis result in cells of abnormal shape and lack of septum synthesis control, probably via different mechanisms (15, 25, 26). Furthermore, proteases, chaperones and secretion proteins, involved in production, folding and/or secretion of cell cycle proteins, may also directly or indirectly affect coordination of cell division and septum formation in *S. aureus*. For example, Clp-protease complexes can target both FtsZ and Sle1, and thereby have major effects on these processes (27-29).

Evidently, cell shape maintenance and control of cell division and septum formation are a complex interplay between many cellular processes where unknown key factors are yet to be discovered. We here identified a hitherto novel protein named SmdA (for <u>s</u>taphylococcal <u>m</u>orphology <u>d</u>eterminant <u>A</u>). We show that the level of the membrane-attached SmdA protein is critical for maintaining normal division progression and morphology in *S. aureus*, as silencing or overexpression of *smdA* resulted in defective cell division and septal cross wall synthesis, as well as increased sensitivity towards cell wall targeting antibiotics.

## Results

### SmdA is a conserved staphylococcal membrane protein

To identify novel proteins potentially involved in cell cycle and morphology control in *S. aureus*, we performed a combined depletion- and subcellular localization analysis of essential staphylococcal proteins with no annotated functions (30-32). From this, we identified a septum-enriched protein (SAOUHSC_01908, named SmdA), whose knockdown resulted in cells of variable sizes that formed large clusters (see below). The function of SmdA in *S. aureus* was investigated further here.

SmdA is a protein of 302 amino acids which is fully conserved in species within the *Staphylococcaceae* family (Fig. S1). The *smdA* gene is monocistronic and located >100 bp away from the neighboring genes (SAOUHSC_01907; unknown function, and *metK*; S-adenosylmethionine synthase). The protein has a predicted N-terminal transmembrane helix and a C-terminal cytoplasmic part with partial homology to a so-called nuclease related domain (NERD) (PF08375, E = 1.08·10^−5^) (Fig. 1A). The domain was named based on distant similarities to endonucleases, and is found in bacterial, archaeal, as well as plant proteins (33). However, the functional role of NERD in bacteria has to our knowledge never been studied. SmdA has been found to be essential in transposon mutagenesis studies in *S. aureus* (30, 32). Growth analysis of SmdA CRISPRi knockdown (hereafter SmdA^down^) in *S. aureus* SH1000 in rich medium at 37°C resulted in a reduction in growth compared to the control strain when spotted on agar plates (Fig. 1B). The growth rate in liquid medium was only slightly affected when CFU and OD_600_ was determined at different time points in the same culture during growth (Fig. S2A). By RT-PCR we verified that the expression of *smdA* was indeed fully knocked down by the CRISPRi system in *S. aureus* SH1000 (Fig. S2B). Since the SmdA^down^ strains were still viable, we attempted to construct deletion mutants of *smdA* by allelic replacement with a spectinomycin resistance cassette using the pMAD-vector (34). However, we were not able to obtain the deletion mutant in *S. aureus* SH1000. Similarly, for *S. aureus* strains NCTC8325-4, HG001 and the MRSA strain COL, SmdA^down^ resulted in reduced growth on agar plates, with HG001 being most affected (Fig. 1B), but we were not able to obtain any deletion mutants in these strains. We therefore used the CRISPRi system in the different *S. aureus* strains to study the phenotypes of SmdA further.

**Fig. 1.**
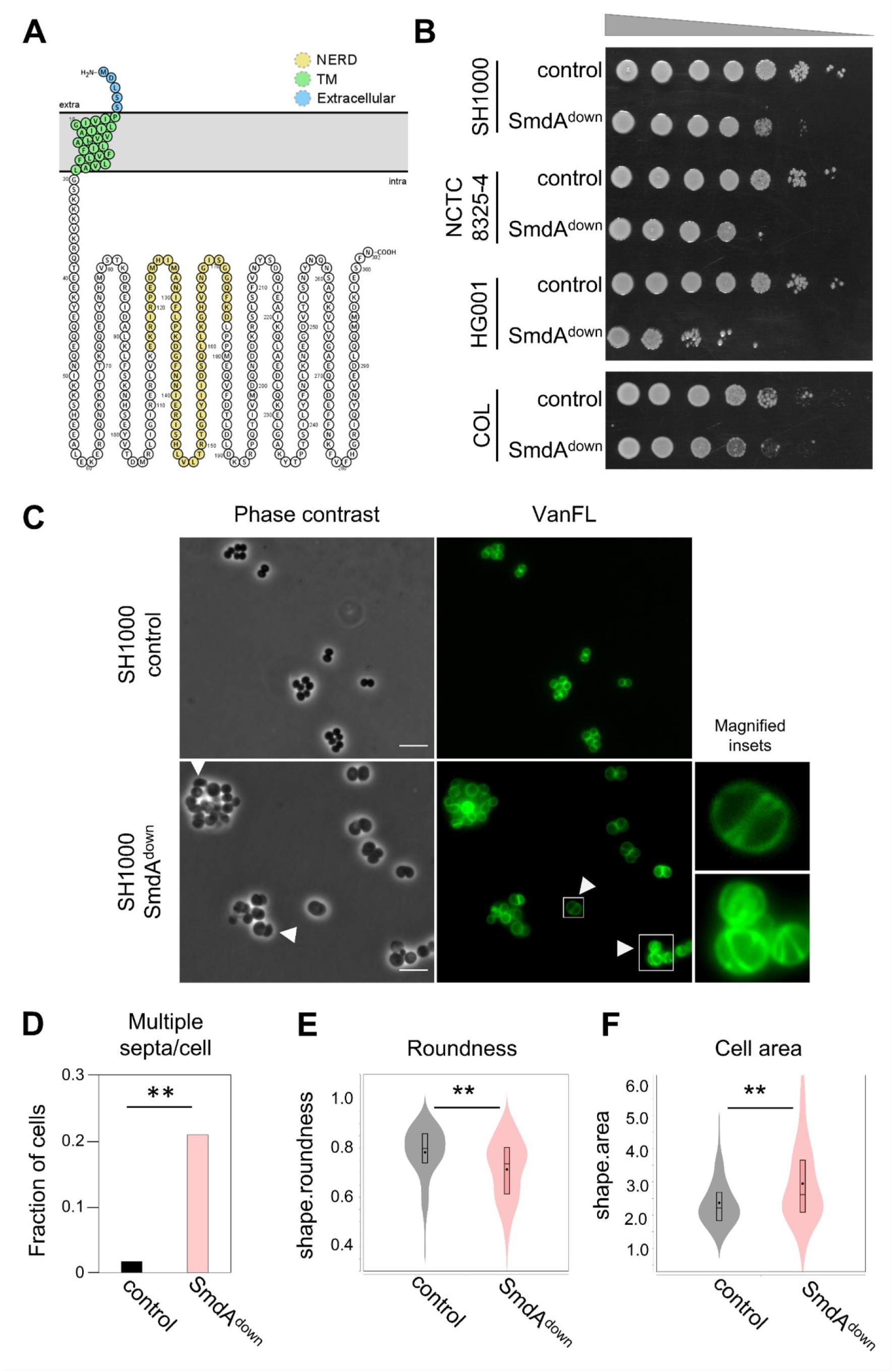
Phenotypes resulting from depletion of SmdA. (**A)** Predicted topology of SmdA using Protter (77). SmdA is predicted to have one transmembrane (TM) helix with a short extracellular N-terminus and a large intracellular domain. The sequence with predicted similarity to the NERD domain is highlighted in yellow. (**B**) Growth on solid media of SmdA knockdown strains (SmdA^down^) in *S. aureus* SH1000 (IM269), NCTC8325-4 (IM311), HG001 (IM312) and COL (IM294). Strains carrying a non-targeting sgRNA were used as controls (IM284, IM307, IM313 and IM295 for the respective strains). From non-induced overnight cultures, 10-fold dilution series were made and spotted onto plates with 300 µM isopropyl-β-D-thiogalactopyranoside (IPTG). (**C**) SmdA^down^ (IM269) and control strain (IM284) analyzed by phase contrast- and fluorescence microscopy of cells stained with the cell wall label VanFL. White arrows point at mis-shaped cells and cells with perturbed septum formation. Magnified insets of representative cells are shown for the VanFL micrographs. Scale bars, 5 µm. (**D**) Fraction of cells with multiple septa per cell for the SmdA^down^ strain IM269 (n = 225) and the non-target control strain IM284 (n = 242) are plotted. The asterisks indicate significant difference (Fisher’s exact test, P<0.001). (**E**) Cell roundness, as determined using MicrobeJ, was used as a measure of the morphology of the cells. Spherical cells will have values close to 1. Cell roundness measures for the control strain IM284 (n = 198) and the SmdA^down^ strain IM269 (n = 191) are plotted. (**F**) Cell area (in µm^2^) as determined using MicrobeJ of the control strain IM284 (n = 198) and the SmdA^down^ strain IM269 (n = 191). In E and F, significant differences between the distributions are indicted by asterisks (**, P < 0.001). P-values were derived from a Mann-Whitney test.

### Depletion and overexpression of SmdA result in cells with highly aberrant cell shapes

During the initial analysis, we observed that SmdA^down^ in *S. aureus* SH1000 resulted in clusters containing cells of variable sizes. The knockdown experiment was repeated, and exponentially growing cells were stained with fluorescent vancomycin (VanFL, binds to non-crosslinked stem peptides throughout the cell wall). SmdA^down^ indeed resulted in severe phenotypic defects (Fig. 1C); increased cell clustering and a large fraction of cells with multiple septa (20.1 % for SmdA^down^ as opposed to 1.6 % for the control strain, Fig. 1D) and abnormal, non-spherical morphology (Fig. 1E). The SmdA^down^ cells were also significantly larger than the control strain (Fig. 1F).

Transmission- and scanning electron microscopy (TEM and SEM) were used to obtain more detailed images of the defects in morphology and septal placement found in SmdA^down^ cells. Strikingly, SmdA depleted *S. aureus* SH1000 displayed highly aberrant septum formation (Fig. 2A). In addition to lysis, cells with several non-perpendicular- or parallel septa were frequently observed, resulting in cells, or small cell clusters, with aberrant morphologies (Fig. 2A). This was also evident from the SEM micrographs, which showed clustered cells with various morphologies (Fig. 2B). Similar phenotypes from TEM and SEM analyses were observed for the NCTC8325-4, HG001 and COL strains, with the HG001 strain being more affected by SmdA^down^ than the other two (Fig. S3 and Fig. S4).

**Fig. 2.**
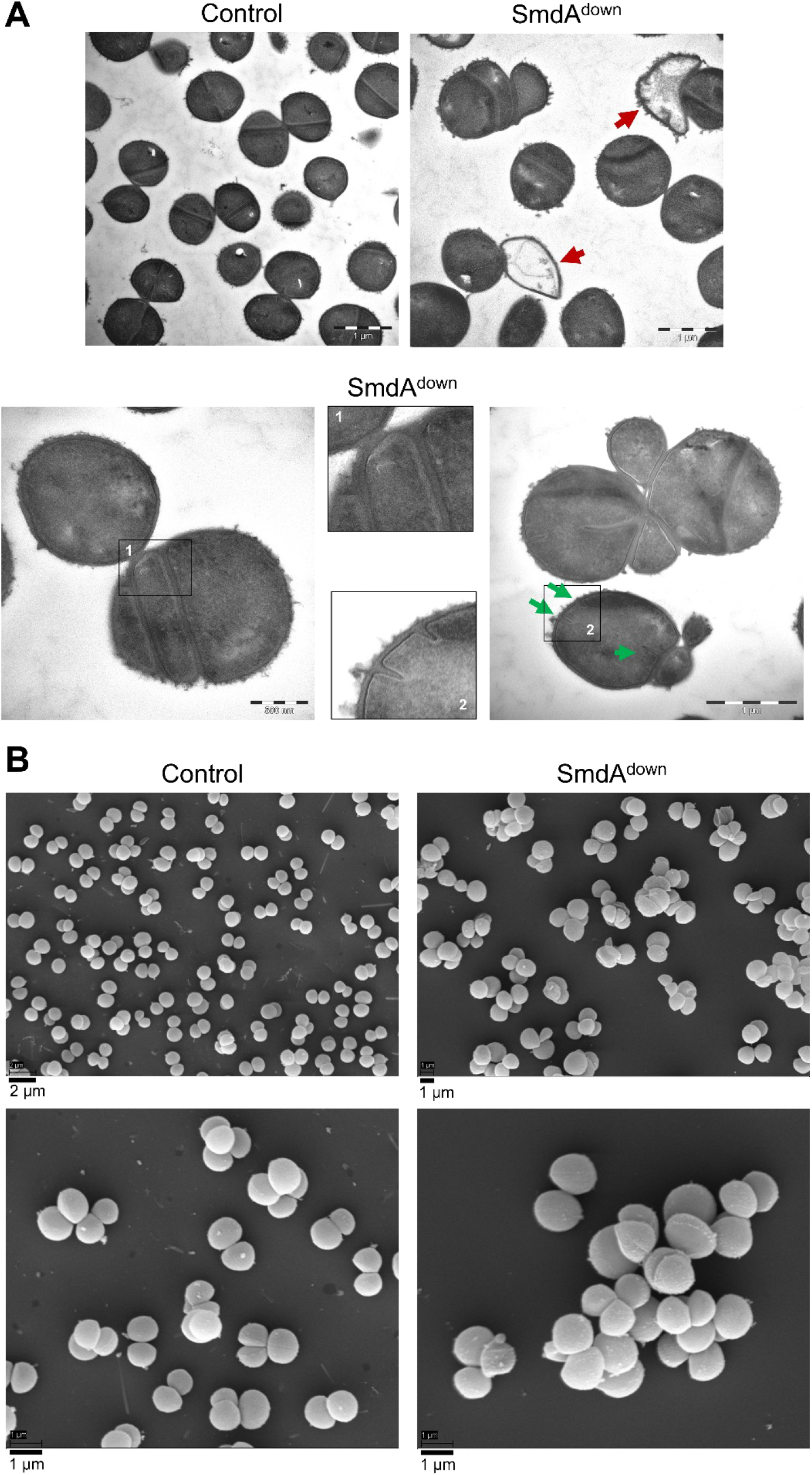
SmdA^down^ in *S. aureus* SH1000 visualized by electron microscopy. (**A**) Transmission electron- and (**B**) scanning electron micrographs of SH1000 CRISPRi control strain (IM284) and SH1000 SmdA^down^ (IM269). In (A), red arrows in the TEM micrographs point at lysed cells. Representative examples of cells with parallel septa or multiple septa are shown. Green arrows point to initiation of septum synthesis at multiple sites within the same cell. Different magnifications are shown, indicated by the scale bars.

Since reduced levels of SmdA led to defects in cell division and morphology of *S. aureus*, we next wondered whether overexpression of SmdA would affect the cells. An ectopic copy of *smdA* under control of an IPTG-inducible promotor in the plasmid pLOW (35) was expressed in *S. aureus* NCTC8325-4. As for SmdA^down^, the cells were stained with VanFL (Fig. S5A). Although less evident than for SmdA^down^, overexpression of SmdA also resulted in clusters of cells with several septa per cell as defined by VanFL staining (Fig. S5A) (3.9 %, n = 181 for SmdA overexpression, compared to 20.1 % for SmdA^down^, and 1.6 % for the control). Together, these knockdown- and overexpression experiments clearly demonstrate that septum formation and splitting in *S. aureus* are dependent on proper levels of SmdA.

### Depletion of SmdA results in increased sensitivity towards antimicrobials targeting cell wall synthesis

Given the negative effect SmdA depletion had on cell morphology and -division, we reasoned that reduced expression of SmdA could influence the sensitivity of *S. aureus* to cell wall targeting antibiotics. To test this, SmdA^down^ strains were treated with PBP-targeting β-lactams (oxacillin, cefotaxime, cefoxitin and imipenem), the glycopeptide vancomycin (blocking cell wall synthesis by targeting the terminal D-Ala-D-Ala on the stem peptides of nascent PG (36)), tunicamycin (targeting TarO and MraY, enzymes involved in the early stages of WTA and PG synthesis, respectively (37)), targocil (targeting the WTA exporter TarG (38)) and Congo Red (inhibitor of the LTA biosynthesis enzyme LtaS (39)) (Table 1). Two antibiotics with alternative targets; tetracycline (targeting protein synthesis) and ciprofloxacin (a quinolone targeting DNA synthesis), were also included.

**Table 1.**
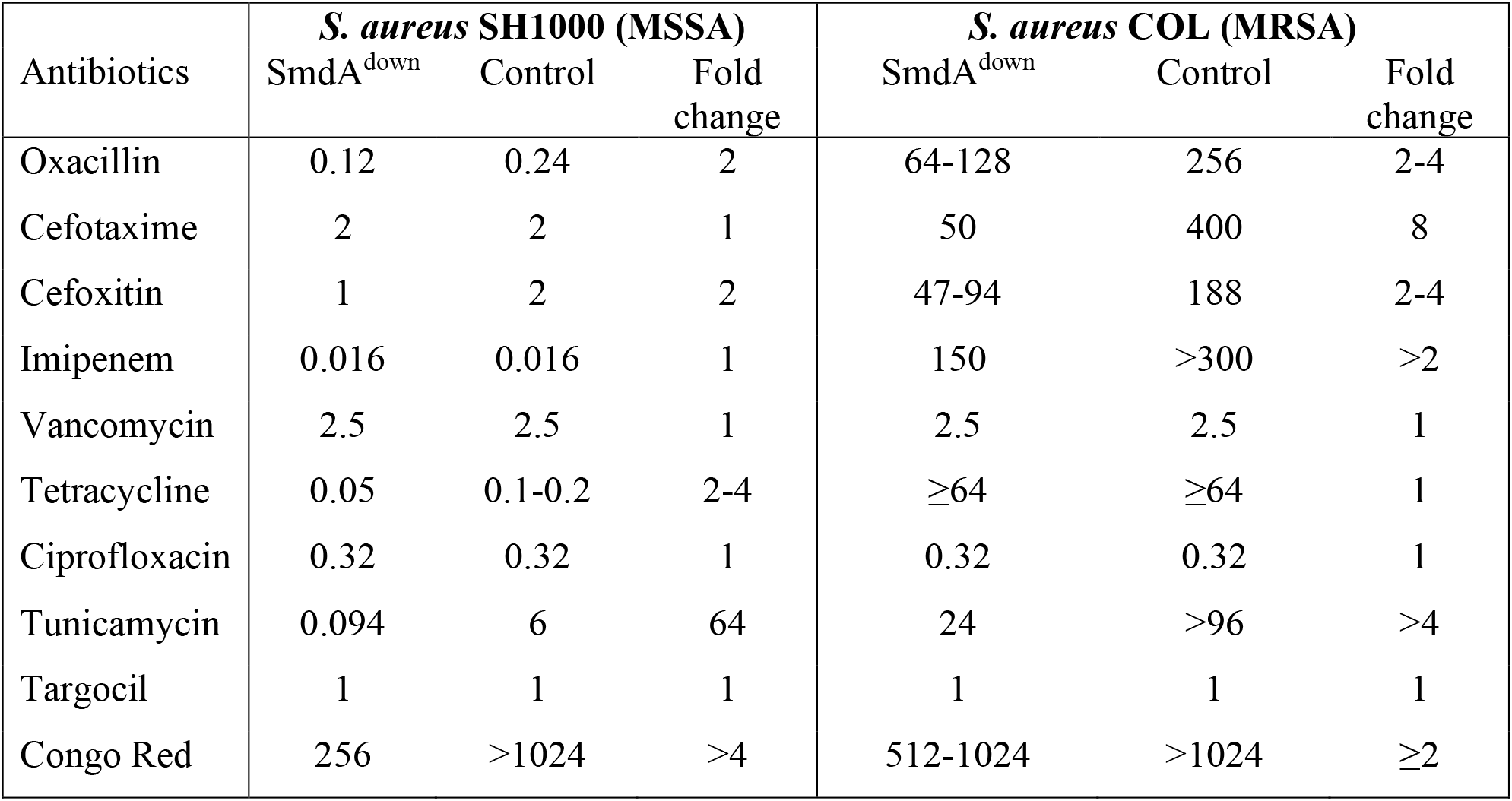
Minimum inhibitory concentration (MIC, in µg/ml) of different antimicrobials when SmdA was depleted in *S. aureus* SH1000 and COL.

For the methicillin-susceptible *S. aureus* (MSSA) strain SH1000, a two-fold reduction in minimum inhibitory concentration (MIC), compared to the control, was observed for oxacillin and cefoxitin in SmdA^down^. More notably, in the MRSA strain COL, SmdA^down^ sensitized the COL strain towards all β-lactams with a 2-to 8-fold reduction in MIC compared to the control (Table 1). SmdA depletion did not seem to significantly influence vancomycin susceptibility. Strikingly, however, we observed that SmdA^down^ cells became highly sensitive towards tunicamycin, with a 64-fold reduction in MIC compared to the control for *S. aureus* SH1000 and >4-fold reduction for COL (40, 41). We therefore also tested the tunicamycin sensitivity of the NCTC8325-4 and HG001 SmdA^down^ strains, which also showed increased susceptibility towards tunicamycin (HG001; 125-fold reduced MIC and NCTC8325-4; 4-fold reduced). While no difference in MIC was observed for targocil, the SmdA^down^ strains also displayed increased sensitivity towards Congo Red. Finally, SmdA depletion in *S. aureus* SH1000 led to a 2-to 4-fold reduction in MIC against tetracycline, compared to the control, however, depletion of SmdA in COL did not change its sensitivity towards tetracycline or ciprofloxacin (Table 1).

### SmdA has no major effects on the TA biosynthetic pathways

The increased sensitivity to tunicamycin and Congo Red in SmdA^down^ strains, prompted us to study whether there were any major alterations in the TA in these cells, although it was also noted that no changes in sensitivity was observed for the WTA export inhibitor targocil (Table 1). Notably, the SmdA^down^ strain displayed morphologies reminiscent of what has previously been reported for cells depleted of TA, i.e., with larger cell sizes, cells with irregular septum formations and reduced splitting (26, 37, 38, 42). It has previously been shown that there is a synthetic lethal relationship between the WTA and LTA biosynthetic pathways (42, 43), and it could therefore be hypothesized that hypersensitivity to tunicamycin could result from deficient LTA biosynthesis in the *smdA*^*down*^ mutants, Using an anti-LTA antibody, we compared the quantity and lengths of LTA in the SmdA^down^ - and control strains for SH1000, NCTC8325-4, HG001 and COL (Fig. S6A). For HG001, the strain with most severe division defects, we observed a reduction in LTA amounts in SmdA^down^ and this can possibly contribute to the severe phenotype of this strain. However, no consistent changes in LTA amounts or lengths were observed between the SmdA depletions and the controls in the four strains. Furthermore, we could not detect any LTA release into the growth medium in the depletion strains, indicating that the stability of LTA (44) was intact (Fig. S6B). We therefore conclude that SmdA does not have any consistent effect on LTA synthesis across strains.

WTA has been shown to protect cells from the LTA-inhibitor Congo Red. Without WTA, cells became hypersensitive towards Congo Red (MIC of <4 µg/ml for *tarO* deletion mutants and >1024 µg/ml for wild-type cells) (45). Although to a much lesser degree, SmdA depletion strains were also more sensitive to Congo Red compared to the controls (Table 1), and we therefore looked into whether WTA could be disturbed in a SmdA^down^ strain. Cells without WTA has previously been shown to lack the dark, electron-dense layer observed in TEM images of crosswalls of *S. aureus* wild type cells (Fig. S6C) (37, 46). TEM images of TarO^down^ and SmdA^down^ showed that TarO^down^, as expected (37, 46), lacked this dark, high-density layer, while it was still present in the SmdA^down^ strain, suggesting that WTA was still produced (Fig. S6C). We also performed Fourier transform infrared spectroscopy (FTIR), which has been used before to detect differences in the composition of WTA due to variable glycosylation patterns (44, 47). As expected, changes in the FTIR spectra were evident in the polysaccharide region (1200 cm^-1^ - 800 cm^-1^) for the TarO^down^ strain compared to the control, with the most significant differences recorded for the peaks at 1076 cm^-1^, 1048 cm^-1^, 1033 cm^- 1^ and 1000 - 970 cm^-1^, representing α- and β-glycosidic bonds in WTA (47). However, no changes were observed between SmdA^down^ and the control (Fig. S6D). Together, these results suggests that SmdA has no major effect on the WTA biosynthetic pathway.

### SmdA is important in several stages of staphylococcal cell division

To identify potential protein interaction partners of SmdA, we next performed a protein pulldown experiment using GFP-trapping with a chromosomal *smdA-m(sf)gfp* fusion strain. Interestingly, the major staphylococcal autolysin Atl, as well as the bifunctional PBP2 were identified, along with 12 other proteins (Table S1). The experiment was repeated in a strain with plasmid-based expression of SmdA-m(sf)GFP, and this setup resulted in an extended list of proteins that were pulled down, probably due to the elevated expression of SmdA-m(sf)GFP. Selecting the proteins for which at least 10 unique peptides were detected and with a fold change of >2 compared to the control, resulted in a list of 57 proteins (Table S1). Several proteins with activity involved in protein folding, secretion and/or degradation (e.g., FtsH, PrsA, SpsB, ClpB, ClpC, SecD) were identified in this assay, in addition to the penicillin-binding proteins PBP1, PBP2, PBP3 and the early division protein EzrA. All the 14 proteins identified in the initial experiment were also pulled down in the second experiment.

Bacterial two-hybrid analyses of the SmdA-PBP1-3 and SmdA-EzrA were performed to see whether these interactions could be reproduced in a heterologous system. The proteins were fused either N- or C-terminally to the domains of adenylate cyclase, an enzyme which catalyzes the production of cyclic adenosine monophosphate (cAMP) and eventually induction of β-galactosidase production when brought in proximity by interaction between the target proteins. Indeed, SmdA interacted with PBP2, as well as PBP1, PBP3 and EzrA in the two-hybrid assays (Fig. 3A). By expressing a version of SmdA without its N-terminal membrane domain (SmdAΔTMH), we also show that the observed interactions between SmdA and the PBPs in this assay occur via the transmembrane segment, while the interaction with EzrA was retained for SmdAΔTMH. The latter suggests that EzrA interacts with the intracellular part of SmdA (Fig. 1A). Furthermore, we also show that SmdA is able to self-interact, and also this interaction is dependent on the transmembrane helix (Fig. S7).

**Fig. 3.**
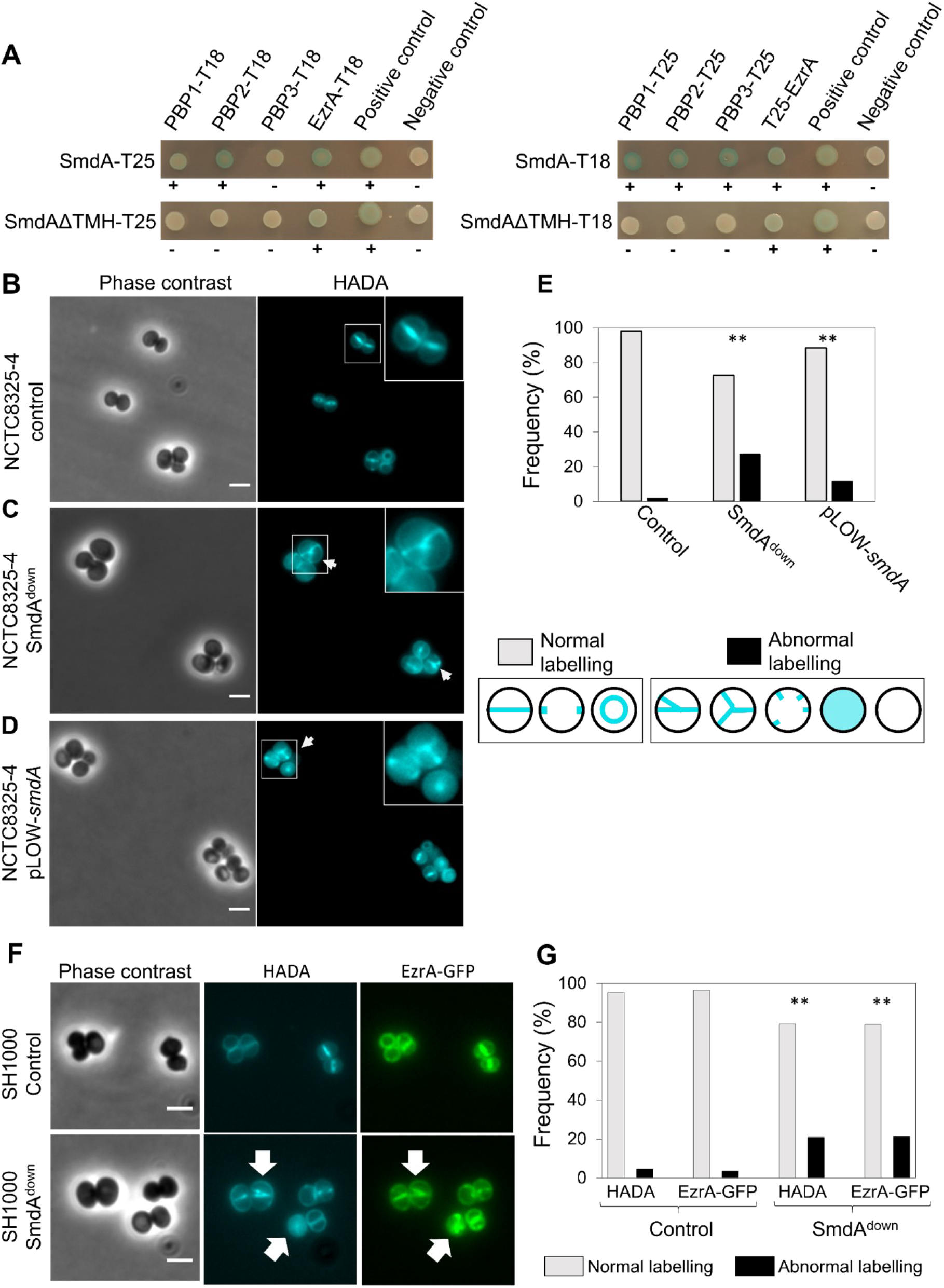
SmdA interacts with important cell division proteins and is necessary for proper localization of EzrA and peptidoglycan synthesis. (**A**) Protein-protein interactions tested with bacterial two-hybrid assays, where SmdA and SmdAΔTMH were tested against PBP1, PBP2, PBP3 and EzrA. The proteins were fused to the T18 or T25 domains as indicated. Blue bacterial spots and plus symbols indicate positive interactions and white spots and minus symbols indicate no interaction. **(B-D)**. Micrographs of HADA labelled *S. aureus* NCTC8325-4 with depletion and overexpression of SmdA. Arrows point at cells with misplaced septum synthesis. Scale bars, 2 µm. **(B)** CRISPRi control strain, IM307 (**C**) SmdA^down^ strain, IM311 and (**D**) SmdA overexpression strain, MK1866. (**E**) Frequency plot of cells with normal or abnormal HADA labelling pattern. Categorization of normal or abnormal labelling patterns are indicated. The number of cells analyzed were 259, 179 and 189 for B, C and D, respectively. The asterisks indicate significant difference from the control (Fisher’s exact test, P<0.001). (**F**) Micrographs showing co-localization of EzrA-GFP and HADA incorporation in *S. aureus* SH1000 strains with (MK1952) or without (MK1953) knockdown of SmdA. Phase contrast- and fluorescence images of HADA labelling and GFP (EzrA-GFP) are shown. Scale bars, 2 µm. Arrows point to cells with abnormal localization of both HADA and EzrA-GFP. (**G**) Frequencies of cells (from F) with normal or abnormal localization of HADA and EzrA-GFP are plotted. The number of cells analyzed were 285 (for MK1953) and 298 (for MK1952). The asterisks indicate significant difference form the respective controls (Fisher’s exact test, P<0.001).

The observed interactions between SmdA and PBPs may suggest that SmdA is somehow important for proper regulation and/or localization of PBPs in *S. aureus* (Fig. 1 and Fig. 2). The VanFL approach used above (Fig. 1C) labels non-crosslinked PG throughout the cell surface. Therefore, to determine more precisely the sites of active PG synthesis in *S. aureus* NCTC8325-4 SmdA^down^ and SmdA overexpression cells, we pulse-labelled the cells for 90 sec with the fluorescent D-amino acid 7-hydroxycoumarincarbonylamino-D-alanine (HADA), a molecule which is integrated into PG by the specific activity of transpeptidases (48). In the control cells, the expected midcell-localized band was observed (Fig. 3B). However, in the SmdA^down^ strain, we observed a more diffuse HADA signal with PG synthesis often occurring at several sites within the cell, forming patterns visible as crosses or Y-shapes (Fig. 3C). Similar observations were made in the cells overexpressing SmdA, although with a lower frequency (Fig. 3D-E). Thus, the site of active transpeptidation (and thus probably at least one of the PBPs) is mis-localized when the levels of SmdA is altered.

Moreover, we also studied whether SmdA could affect the localization of EzrA by knocking down *smdA* in a strain expressing a chromosomal *ezrA-gfp* fusion (Fig. 3F). Indeed, abnormal localization patterns for EzrA-GFP was observed with similar frequency as the abnormal HADA labelling (Fig. 3G). These results thus suggest that depletion of SmdA, directly or indirectly, influences the localization of both early and late divisome proteins in *S. aureus*.

Furthermore, the major autolysin Atl was also pulled down with SmdA. Atl is a secreted multidomain enzyme, which is processed to an acetylmuramyl-L-alanine amidase and β-N-acetylglucosaminidase, involved in septal cross wall splitting resulting in daughter cell separation (19). SmdA^down^ cells frequently displayed increased clustering (Fig. 2, Fig. S4). This indicates reduced cross wall splitting, a phenotype also observed in Δ*atl* mutants and WalKR depleted cells (22, 49). WalKR is the two-component regulatory system controlling the expression of *atl* and other cell wall hydrolase encoding genes (22). To assess the reduced cross wall splitting phenotype in more detail, we performed Triton X-100-induced autolysis assays on the cultures. Indeed, reduced autolysis was observed in SmdA^down^ cells, demonstrating reduced autolytic activity (Fig. 4A). The lysostaphin sensitivity was also reduced (Fig. 4B), suggesting alterations in the cell wall affecting the lytic properties of this enzyme. It should be noted that the resistance towards Triton X-100- and lysostaphin-induced autolysis in the SmdA^down^ strain was clearly less compared to the control strain where *walR*, and thus all the regulated autolysins, was knocked down. It should also be noted that Triton-X induced autolysis was not severely altered upon overexpression of SmdA (Fig. S5B).

**Fig. 4.**
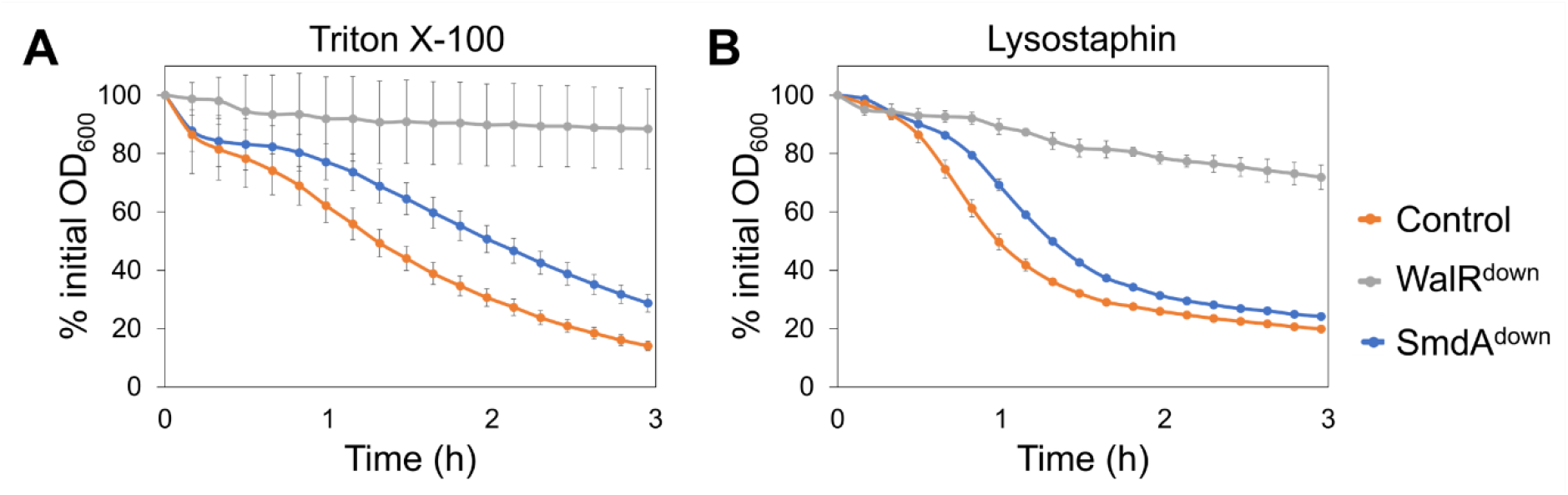
Autolysis of *S. aureus* SH1000 depletion strains (control = IM284; WalR^down^ = IM293; SmdA^down^ = IM269) measured in presence of (**A**) 0.5 % Triton X-100 and (**B**) 100 ng/ml lysostaphin. Results presented as % of initial OD_600_. Error bars represent standard error calculated from four technical replicates.

### SmdA localizes at the septal region after FtsZ

The SmdA-m(sf)GFP fusion protein displayed a septum-enriched signal when expressed ectopically from the low copy-number plasmid pLOW (Fig. 5A). In order to analyze the SmdA localization with native expression levels, a chromosomally integrated version of *smdA-m(sf)gfp* was made, in which SmdA was expressed with m(sf)GFP fused to its C-terminus. Localization analyses in these cells further confirmed the septum-enriched localization of the fusion protein (Fig. 5A), with an average septum/periphery fluorescence signal ratio of 3.4 (n = 52, see Materials and Methods). To visualize the localization of SmdA relative to the cell division process, we used FtsZ as a marker. Expression of *ftsZ*-fusion genes in the *smdA-m(sf)gfp* chromosomal fusion strain, however, only resulted in strains with extremely poor growth, suggesting that the cells did not tolerate such double-labelling. Instead, we therefore created a double-labelled strain in which SmdA-mYFP and FtsZ-mKate2 were co-expressed from plasmids, while the native *smdA* and *ftsZ* genes were still present on the chromosome. Structured illumination microscopy (SIM) analysis showed that SmdA-mYFP is localized around in the membrane when the Z-ring is formed (Fig. 5B), and that a septum-enriched localization occurs as the septal cross wall is being synthesized. This was further confirmed by stimulated emission depletion (STED) microscopy analysis (Fig. 5C). STED imaging also revealed that there was no apparent co-localization of SmdA-mYFP and FtsZ-mKate2 in newborn cells before FtsZ-constriction initiates. Combined, the localization of SmdA is reminiscent of the localization of PBPs in *S. aureus* (11), but appears to localize to the septal area after the early divisome proteins such as EzrA (50, 51).

**Fig. 5.**
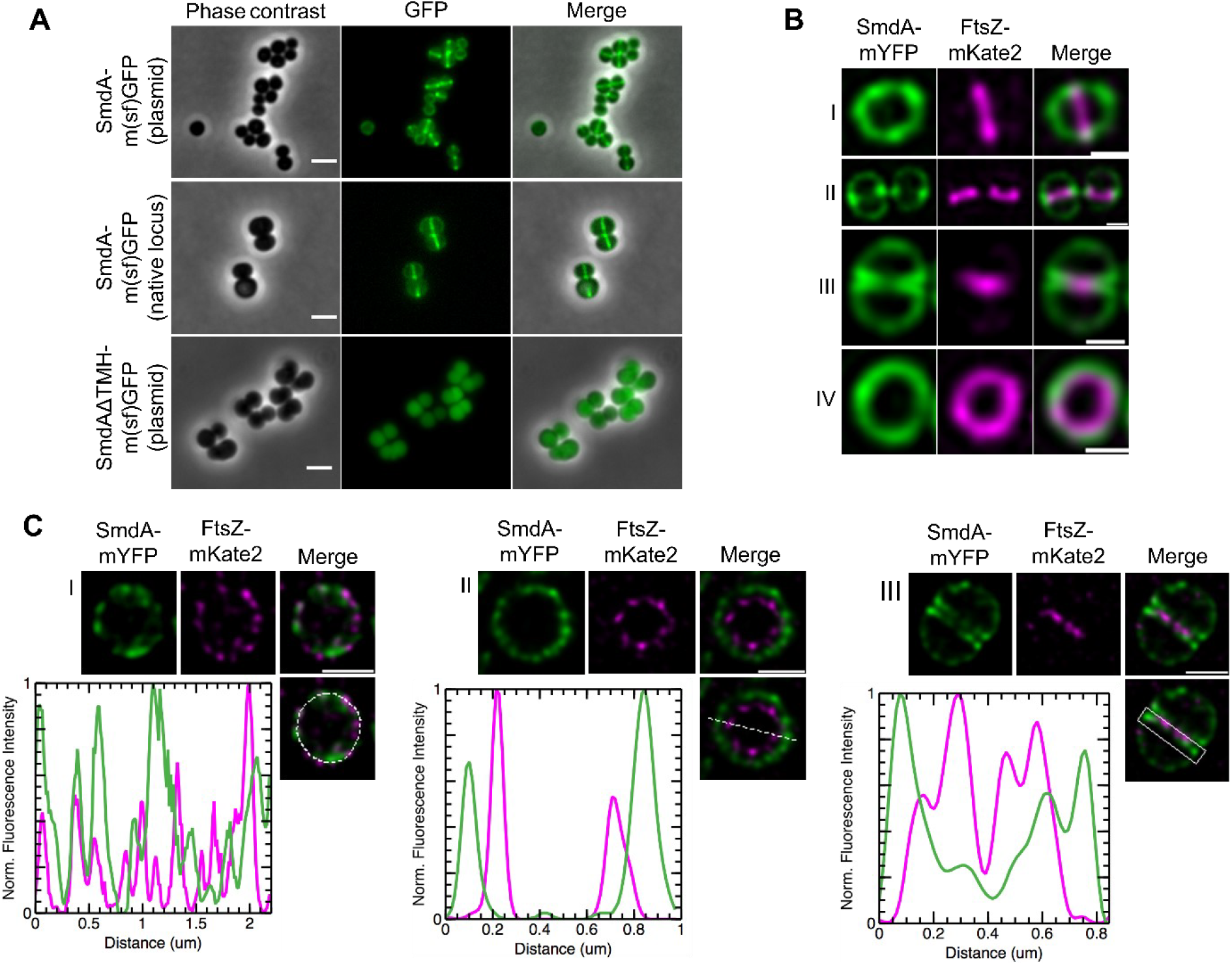
Subcellular localization analysis of SmdA. (**A**) Micrographs of cells with induced expression of SmdA-m(sf)GFP from plasmid (top panel, IM104), native chromosomal expression of the SmdA-m(sf)GFP fusion protein (middle panel, IM308), and expression of SmdAΔTMH-m(sf)GFP from plasmid (bottom panel, IM373). Scale bars, 2 µm. (**B**) SIM images of fixed *S. aureus* SH1000 with plasmid-expressed SmdA-mYFP and FtsZ-mKate2 (HC060). (**B-I**) Side-view of a cell showing that FtsZ localizes at septum before arrival of SmdA. As cell division progresses, (**B-II**) SmdA concentrates at sites where septum formation is initiated, and (**B-III**) displays a septal localization at the two septal membranes as FtsZ constricts and septum formation proceeds. (**B*-*IV**) Top-view of a cell showing the FtsZ-ring inside of the SmdA-ring. All scale bars, 0.5 µm. (**C**) STED images of fixed *S. aureus* SH1000 with plasmid-expressed SmdA-mYFP and FtsZ-mKate2 (HC060), where line scans show fluorescence intensity of selected areas. (**C-I**) At an early stage of cell division, the rings of both SmdA and FtsZ have a similar diameter, but do not overlap in their distribution, and a heterogenous distribution with a patchy signal is observed. (**C-II**) Top-view of a cell as cell division progresses, with the FtsZ-ring laying inside of SmdA, approximately 100-150 nm apart from each other. (**C-III**) Side-view of a cell showing how FtsZ is located innermost, and how SmdA is located at the two septal membranes. Scale bars, 0.5 µm.

The SmdA protein is anchored to the membrane by a single transmembrane helix (Fig. 1A). To verify its significance for the septal localization, we ectopically expressed a version of SmdA without the transmembrane helix (SmdAΔTMH). The N-terminally truncated SmdAΔTMH-m(sf)GFP localized to the cytoplasm of the cells (Fig. 5A). This shows that while the transmembrane helix is critical for protein-protein interactions and subcellular localization of SmdA, the interaction observed between EzrA and SmdAΔTMH (and full-length SmdA) (Fig. 3A) is probably not involved in determining the localization of this protein.

During these latter experiments, we noticed that SmdAΔTMH-m(sf)GFP overexpression led to cells with obvious morphology defects. In fact, overexpression of SmdAΔTMH (without the GFP-tag) (Fig. 6A) strikingly resulted in a more extreme phenotype than overexpression of full length SmdA (Fig. 3D, Fig. S5). The cells were often inflated or with a bean-shaped appearance (Fig. 6A). HADA pulse labelling of these cells revealed a highly abnormal PG incorporation pattern, with the fluorescent signal forming both condensed clumps and Y-shapes in different directions in almost 50 % of the cells (Fig. 6A and B). This demonstrates that membrane attachment is critical for the localization and function of SmdA.

**Fig. 6.**
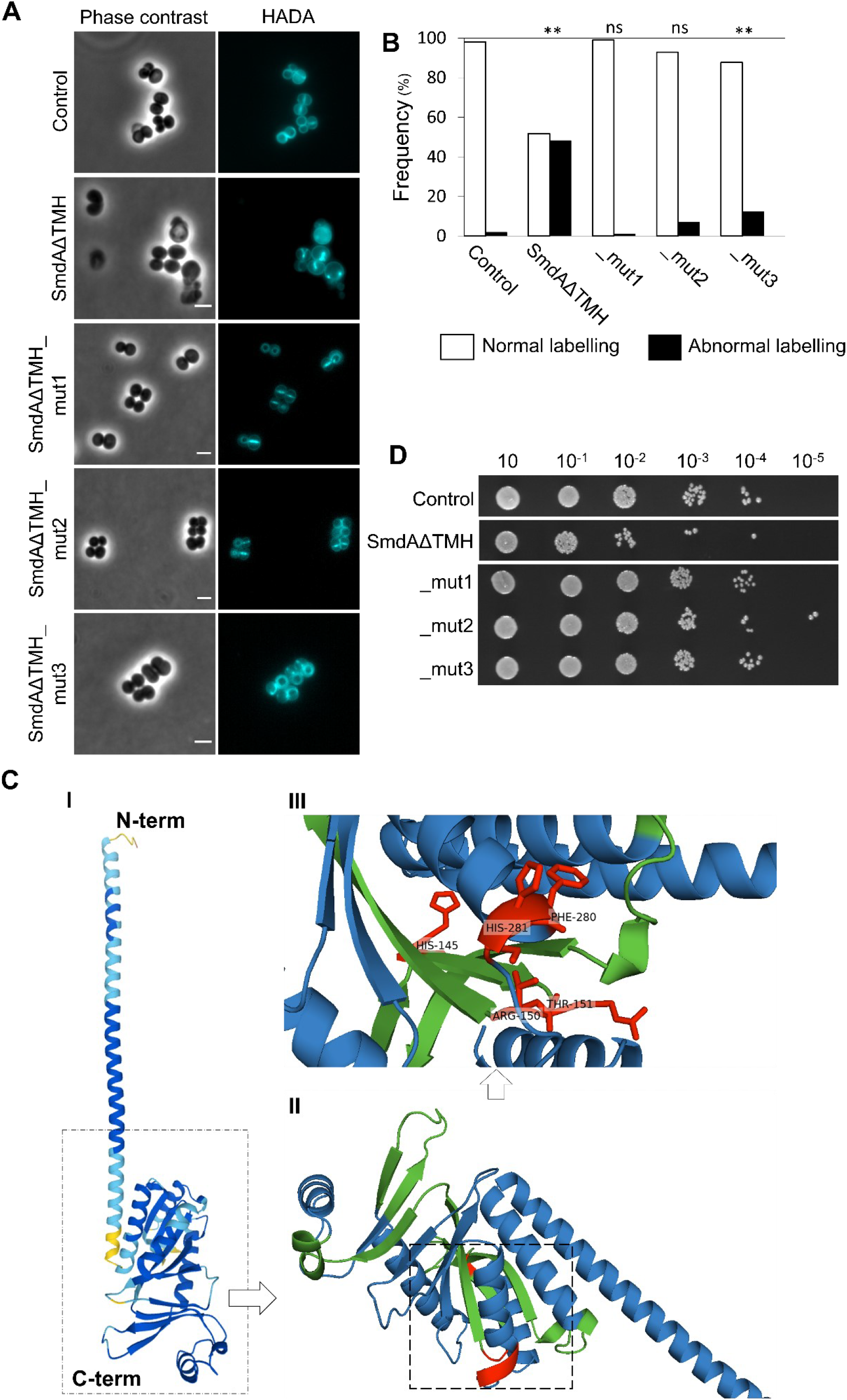
Membrane attachment and the NERD domain are important for SmdA function. HADA stained *S. aureus* NCTC8325-4 control cells not expressing any SmdA-variant (IM307) and cells overexpressing SmdAΔTMH (MK1911), SmdAΔTMH_mut1 (IM377), _mut2 (IM378) and _mut3 (IM379). Scale bars, 2 µm. (**B**) The frequency of cells from panel A with normal or abnormal HADA staining is plotted. The number of cells plotted were 259 the control, 82 for SmdAΔTMH, 101 for _mut1, 113 for _mut2 and 122 for _mut3. The asterisks indicate significant difference form the control (Fisher’s exact test, P<0.001). ns indicates that there is not significant difference from the control. (**C**) Structure of SmdA, predicted by AlphaFold (52), where (**C-I**) shows the structure colored by a “per-residue confidence score” (pLDDT) with dark blue pLDDT > 90, light blue 90 > pLDDT > 70, and yellow 70 > pLDDT > 50, (**C-II**) shows the structured domain magnified with the predicted NERD colored in green and residues that have been mutated in red. (**C-III**) A magnified inset of (II) with annotated residues. The positions of the mutated residues are indicated in red. (**D**) Growth assay on solid media of *S. aureus* NCTC8325-4 expressing SmdAΔTMH, SmdAΔTMH_mut1, _mut2 and _mut3.

### Conserved amino acids in the predicted NERD domain are critical for the function of SmdA

The functional importance of the NERD domain (Fig. 1A) has, to our knowledge, neither been studied nor verified previously in any bacterial protein. As mentioned, we observed that overexpression of SmdAΔTMH led to severe phenotypic changes in the staphylococcal morphology (Fig. 6A). We decided to use this as a tool to gain insight into whether the predicted NERD domain is important for the function of SmdA. Multiple sequence alignment of SmdA proteins from different staphylococcal species (Fig. S1) revealed conserved residues in this domain, and site directed mutagenesis was used to create two versions of SmdAΔTMH containing mutations in the NERD domain: one in which H145 was changed to Ala (mut1), and a second containing the mutations R150A and T151A (mut2). Furthermore, we also observed from the alignment that the C-terminus of SmdA is highly conserved (Fig. S1), and another C-terminally mutated variant was created (mut3; F280A, H281A). The three-dimensional structure of SmdA, predicted by AlphaFold (52), suggests that the N-terminal part of the protein folds into a long α-helix and that the NERD domain is located within the C-terminal structured part of the protein (Fig. 6C-II). The mutated SmdAΔTMH versions were overexpressed in a wild-type background. Expression of these mutated variants indeed partially, or fully, abolished the phenotypic defects observed when overexpressing the non-mutated SmdAΔTMH version. HADA staining of mut1 (H145A) showed septum placement and PG synthesis comparable to wild-type cells, while mut2 (R150A, T151A) and mut3 (F280A, H281A) also clearly reduced the functionality of the protein (Fig. 6A and B). Furthermore, the growth inhibition upon overexpression of SmdAΔTMH (Fig. 6D) was also abolished when the mutated variants were expressed under the same conditions. This strongly suggests that the given residues in both the predicted NERD and the conserved C-terminus are important for the function of SmdA.

## Discussion

The results presented here provide the first functional insights into the conserved staphylococcal protein SAOUHSC_01908, here named SmdA. SmdA, a protein conserved within the *Staphylococcaceae* family, is critical for maintaining cell morphology and for proper progression of cell division in different *S. aureus* strains (including MRSA and MSSA). Unbalanced levels, as well as loss of proper subcellular localization, of SmdA, result in severe cell morphology defects due to mis-localized cell division, uncontrolled cell wall synthesis, and lack of proper cross wall splitting.

We observed that knockdown of SmdA increased the sensitivity of *S. aureus* to several cell wall targeting antibiotics, including β-lactams and tunicamycin (Table 1). Many of the frequently used antibiotics, including β-lactams, target cell wall synthesis, but the rise of both methicillin-resistant- and vancomycin-resistant *S. aureus* (MRSA and VRSA) have made *S. aureus* infections more difficult to treat and novel anti-staphylococcal targets and strategies are needed. Targeting cell cycle proteins have been shown to re-sensitize resistant staphylococci towards existing antibiotics due to synergistic effects (53-55). For example, inactivation of some proteins involved in WTA synthesis sensitizes MRSA to β-lactams (37, 56). The same is also the case for a diversity of factors contributing to cell division and cell wall biogenesis, including FtsZ, FtsA, PBP4 and most proteins involved in the PG synthesis pathway (55). The secretion-associated proteins SecDF and the chaperones PrsA and HtrA1 have also been tightly linked to β-lactam sensitivity (57, 58), and recently, inactivation of the autolytic cell wall amidase Sle1 (21), or the membrane proteins AuxA and AuxB (44), were shown to result in increased sensitivity to β-lactams. Mechanisms of re-sensitization in these cases vary and may be a result of weakened cell wall, as well as inactivation or mis-localization of PG synthesis, including the key resistance determinant PBP2A. The large number of factors affecting β-lactam sensitivity reflects the tight links between different processes involved in cell wall biosynthesis, and most likely SmdA affects several steps in the cell division process (Fig. 1-3).

The exact mechanism by which SmdA affects these processes remains to be determined. Our results suggest that cell division and PG synthesis, but not teichoic acid biosynthesis, is affected. This is supported by observations showing mis-localization of cell division and PG synthesis in SmdA^down^ cells (Fig. 1-3). Furthermore, SmdA^down^ cells displayed hypersensitivity towards tunicamycin (targeting enzymes involved both in PG and WTA synthesis), but not towards targocil (targeting the WTA exporter only) (38). Possibly, SmdA influences the localization of cell division and cell wall synthesis by direct protein-protein interactions. This is supported by the SmdA-EzrA and SmdA-PBP1-3 interactions, combined with the abnormal localization of EzrA-GFP and HADA incorporation in SmdA^down^ cells (Fig. 3). In fact, the division defects observed in SmdA^down^ cells (Fig. 1-3) are reminiscent of previous studies of *ezrA* null mutants, strains with point mutations in FtsZ or cells exposed to Z-ring inhibitors (50, 51, 59, 60). However, the contribution of the identified interactions to the observed phenotypes must be investigated further, particularly since the subcellular localization of SmdA-m(sf)GFP does not appear to overlap with the Z-ring throughout the cell cycle. The SmdA-localization, which depends on the transmembrane helix, is instead more similar to the reported localization of PBPs (Fig. 5) (11, 50, 51).

Lack of cross wall splitting is another prominent feature of the SmdA depleted cells (Fig. 2 and 4). After septum formation, cells remain attached, forming clusters of mis-shaped cells. Indeed, the autolytic activity was reduced in the SmdA^down^ cells (Fig. 4). The major autolysin Atl was also pulled down together with SmdA-m(sf)GFP as one of the major hits (Table S1). Atl is an extracellular processed protein whose amidase and glucosaminidase domains together process PG in the septum to allow splitting. It is possible that SmdA somehow influences the export or processing of Atl. In this context, it is interesting to note that proteases, foldases and chaperones (FtsH, PrsA, SpsB, ClpC, ClpB, HtrA1) were also pulled down with SmdA (Table S1). It remains to be determined whether these interactions have any functional relevance, however, perturbation of such pathways may have major consequences for cell division and septal placement as they are important for proper folding and secretion of key proteins involved in such processes. This has, for example, previously been demonstrated for the chaperone ClpX, which is critical for coordination of autolysins and cell division proteins (29). Involvement in such a mechanism could also explain the pleiotropic phenotypes observed in SmdA^down^ cells, affecting several stages in the cell division process.

A part of SmdA displays limited similarity to the so-called nuclease related domain (NERD). This domain, which is found in a broad range of bacterial species and in some archaea and plants, was initially identified on a virulence-plasmid from *Bacillus anthracis* (33), and has later been suggested to belong to a superfamily of phosphodiesterases (61). The majority of NERD proteins are single-domain proteins, but they are also occasionally found together with a kinase-, pseudokinase- or helicase domain (62). The actual function of the NERD has, to our knowledge, however, not been studied experimentally. The results in this paper show that conserved residues in the predicted NERD are important for the functionality of SmdA, thus demonstrating for the first time a functional role of this domain (Fig. 6). Although highly speculative at this point, it would be interesting if a cell division factor such as SmdA had DNA-interacting capabilities and thus forming a potential link between the membrane-associated cell division proteins and the nucleoid. It should also be noted that several of the most conserved residues of the NERD domain (33) are not found in SmdA, suggesting a functional diversity among proteins harboring homology to this domain and implying that the NERD of SmdA may have functions unrelated to the nucleoid.

It is also interesting to note that the effect of SmdA knockdown was different between strains. For example, SmdA^down^ resulted in a more severe phenotype in HG001 compared to NCTC8325-4 and SH1000 (Fig. 1, Fig. S3-S4), although these strains derive from the same parent (63). While HG001 contains prophages, NCTC8325-4 and SH1000 have been cured of these, and it could for example be speculated that the stress imposed by knockdown of SmdA somehow results in prophage induction in HG001 and thereby a more severe phenotype. Understanding of these and other strain differences could also help to further pinpoint the exact function of SmdA.

The staphylococcal genome encodes hundreds of essential proteins, which all represent potential target sites for antimicrobials. To be able to fully exploit the antibiotic target repertoire, it is critical to understand how essential proteins are involved and linked between different cell cycle processes. In this work, we identified and characterized SmdA as a novel factor essential for cell morphology and cell division in *S. aureus*. Based on the results presented here, future research should aim at pinpointing the molecular mechanism by which SmdA affect different stages of the cell division process. Finally, since SmdA depletion results in increased sensitivity to β-lactams, it may as such also be a possible future target for combatting β-lactam resistance by re-sensitizing MRSA to these antibiotics.

## Materials and Methods

### Bacterial strains, growth conditions and transformations

*Escherichia coli* IM08B (64), *E. coli* XL1-Blue and *E. coli* BTH101 were grown in lysogeny broth (LB) at 30-37°C with shaking or on lysogeny agar (LA) plates at 30-37°C, with 100 µg/ml ampicillin and/or 50 µg/ml kanamycin for selection. *S. aureus* SH1000, NCTC8325-4, HG001 and COL were grown in brain-heart infusion (BHI) broth or tryptic soy broth (TSB) at 37°C with shaking or on BHI/TSA plates at 37°C. For selection, 10 µg/ml of chloramphenicol, 5 µg/ml of erythromycin, 15 µg/ml neomycin or 100 µg/ml spectinomycin were added. For induction of gene expression, 50 or 300 µM isopropyl β-D-1-thiogalactopyranoside (IPTG) or 2.5 ng/ml anhydrotetracycline (ATc) were added to the bacterial cultures.

A standard heat shock protocol was used for transformation of *E. coli* IM08B. Isolated plasmids from *E. coli* were transformed into *S. aureus* by electroporation. Preparation of electrocompetent *S. aureus* cells and electroporation were performed according to Löfblom et al (65). Bacterial strains used in this study are listed in Table S2.

### Genetic modifications

For all cloning, DNA fragments were amplified from *S. aureus* SH1000 genomic DNA, and cloning was performed with restriction digestion (New England Biolabs [NEB]) and subsequent ligation using T4 DNA Ligase (NEB), unless otherwise stated. Ligation mixtures were transformed into *E. coli* IM08B, and sequence-verified plasmids were transformed into *S. aureus*. All strains and primers used in this study are listed in Table S2 and Table S3, respectively.

### Fluorescent fusion constructs

#### Construction of pLOW-smdA-m(sf)gfp

Monomeric superfolder GFP, *m(sf)gfp*, was initially fused to *ftsZ* in the plasmid pLOW (35). *m(sf)gfp* was amplified from the plasmid pMK17 (66) using forward primer im1_linker_FP_F_BamHI (annealing to the linker sequence) and reverse primer im2_m(sf)gfp_R_NotI_EcoRI. The amplified fragment encodes a linker sequence N-terminally of the *m(sf)gfp* gene. The fragment was digested with BamHI and EcoRI and subsequently ligated into the corresponding sites of the plasmid pLOW-*ftsZ-gfp* to replace the original *gfp*-gene with the linker_m(sf)gfp sequence. SAOUSHSC_01908 was then amplified with primers im77_SA1908_F_SalI_RBS and im78_SA1908_R_BamHI, the product was digested with SalI and BamHI and ligated into the corresponding sites of pLOW-*ftsZ-m(sf)gfp* to create pLOW-*smdA-m(sf)gfp*.

#### Construction of pMAD-smdA-m(sf)gfp_aad9 for chromosomal integration

To tag the chromosomal *smdA*, the plasmid pMAD-*smdA-flag_aad9* was constructed initially. The insert in this plasmid (*smdA-flag_aad9*) was assembled by overlap extension PCR (primers im147-im152), with the flag-tag sequence inserted by primer overhangs, and cloned into pMAD (34) using restriction enzymes MluI and BamHI and T4 DNA Ligase (NEB). The insert was designed so that the flag-tag could be removed using NotI and SpeI. The *m(sf)gfp* sequence was amplified from template pMK17 (66) using the primers im153 and im154. The PCR product was digested with NotI and SpeI and subsequently ligated with NotI- and SpeI-digested pMAD-*smdA-flag_aad9*. The ligation mix was transformed to *E. coli* IM08B, and the resulting plasmid, pMAD-*smdA-m(sf)gfp_aad9*, was verified by sequencing and transformed into electrocompetent *S. aureus* SH1000. Integration of *smdA-m(sf)gfp* in the native locus of *smdA* using the temperature-sensitive pMAD-system was carried out as previously described (34).

#### Construction of pHC-ftsZ-mKate2

*ftsZ-mKate2* was amplified from the plasmid pLOW-*ftsZ-mKate2*. pLOW-*ftsZ-mKate2* was made in a similar manner as described for pLOW-*ftsZ-m(sf)gfp*, but with reverse primer im5_mKate_R_NotI_EcoRI and genomic DNA derived from strain MK119 (67) as template for amplification of *mKate2*. Then, *ftsZ-mKate2* was amplified with primers USHC109 and USHC148, generating a product with SalI and MluI as overhang. The PCR product and the plasmid pSK9065 (68) were digested with SalI and MluI and thereafter ligated.

#### Construction of pLOW-smdA-Myfp

This plasmid was constructed by using pLOW-*ftsZ-mYFP* as starting point, made similarly as described for pLOW-*ftsZ-m(sf)gfp*, but with reverse primer im3_cfp_myfp_R_NotI_EcoRI and plasmid pMK20 (lab collection) as template for amplifying *mYFP*. The plasmids pLOW-*smdA-m(sf)gfp* (described above) and pLOW-*ftsZ-mYFP* were digested with SalI and BamHI and subsequently ligated, resulting in pLOW-*smdA-mYFP*.

### CRISPRi constructs

#### Construction of pCG248-sgRNA(smdA)

For gene knockdowns, the CRISPR interference (CRISPRi) system developed by Stamsås et al. (69) was used. In this system, *dcas9* is placed downstream of an IPTG-inducible promoter in the plasmid pLOW-*dcas9*. A second plasmid is carrying the single guide RNA (pCG248-sgRNA(*x*), where *x* represents the targeted gene), which constitutively expresses the sgRNA, including the 20 nt base-pairing region specific for the gene to be knocked down. The pLOW-*dcas9* plasmid contains an erythromycin resistance gene, and the pCG248-sgRNA(*x*) plasmid a chloramphenicol resistance gene. The gene-specific 20 nt sequences were replaced in the pCG248-sgRNA(*x*) plasmids using an inverse PCR approach as described earlier (69) using primers mk299 and mk323.

#### Construction of pLOW-dCas9_aad9

In order to make the CRISPRi system compatible with the MRSA strain COL (which is intrinsically erythromycin resistant), the erythromycin resistance gene *ermC* in pLOW-*dcas9* was replaced with *aad9*, encoding a spectinomycin resistance cassette. The primers im183 and im184 were used to amplify the entire pLOW-*dcas9* plasmid, except the *ermC* gene. The *aad9* gene was amplified with primers im185 and im186 using pCN55 (70) as template, where im185 contained the sequence of im183 as overhang, and im186 the sequence of im184 as overhang. Thus, the two fragments had overlapping sequences and were fused using NEBuilder® HiFi DNA Assembly (NEB). The construct was transformed to *E. coli* IM08B, and the plasmid verified by PCR and sequencing. For CRISPRi in *S. aureus* COL, the pLOW-*dcas9_aad9* plasmid was used along with pCG248-sgRNA(*x*).

### Construction of plasmids used for overexpression studies

#### Construction of pLOW-smdA and mutated versions

*smdA* was amplified from *S. aureus* genomic DNA using primers im77_SA1908_F_SalI_RBS and mk517_1908_R_NotI. The fragment was digested with SalI and NotI and ligated into the corresponding sites of plasmid pLOW-*dcas9* (69) to produce the plasmid pLOW-*smdA*, with IPTG-inducible overexpression of *smdA*. pLOW-*smdAΔTMH* was constructed in a similar manner, except that primer mk518_1908_F_RBS_SalI was used instead of im77 to remove the 29 N-terminal amino acids of SmdA, predicted to encode the TMH and extracellular part. Site-directed mutagenesis in the two plasmids was performed by a two-step overlap extension PCR approach, where the mutations were introduced in the primers. The primers mk519 and mk520 were used for introducing mutation H145A (mut1), mut2 (R150A, R151A) was made with primers mk521 and mk522, and mut3 (F280A, H281A) with primers mk529 and mk530.

### Phase contrast- and fluorescence microscopy analysis

For induction of plasmid-encoded fluorescent fusions, exponentially growing cultures were diluted to OD_600_ 0.05 in medium with 50 µM IPTG an incubated for 2 hours prior to microscopy. For CRISPRi-knockdown and overexpression experiments, cultures of OD_600_ =0.4 were diluted 250-fold in medium with 300 µM IPTG and grown until OD_600_ was approximately 0.4. For labelling of cell wall, BODIPY™ FL vancomycin (VanFL) (Invitrogen) was added. To label newly synthesized peptidoglycan, HADA (van Nieuwenhze group, Indiana University) was added actively growing cultures (OD_600_ approximately 0.4) at a final concentration of 250 µM. The cells were incubated with HADA at 37°C for 90 sec and then put on ice. The cultures were pelleted by centrifugation at 10 000 × *g* at 4°C for 1 min, washed with cold 1 × PBS, pH 7.4 and finally resuspended in 25 µl PBS. Cells were subsequently added to agarose pads and microscopy was performed on a Zeiss AxioObserver with ZEN Blue software. An ORCAFlash4.0 V2 Digital complementary metal-oxide semiconductor (CMOS) camera (Hamamatsu Photonics) was used to capture images through a 100 × PC objective. HPX 120 Illuminator (Zeiss) was used as a light source for fluorescence microscopy.

All microscopy analyses were repeated at least two times. Images were processed and prepared for publication using Fiji (71). For analysis of cell roundness, cell area and fluorescent labelling patterns, MicrobeJ (72) was used to determine cell outlines. Outlines were corrected manually, when necessary. Cell area and roundness as a measure of morphology were calculated and plotted using MicrobeJ. After cell outline detection, categorization of cells into normal or abnormal labelling patterns were done manually using images from independent experiments. The septum/periphery fluorescence signal ratio was measured as described before (73) in cells with full septal signal of SmdA-GFP. A fluoresce ratio between 2.5 and 3.5 is typical for septum-enriched proteins (11).

### Structured illumination microscopy (SIM) and stimulated emission depletion (STED) microscopy

Super-resolution SIM imaging was performed using a Zeiss ELYRA PS.1 microscope equipped with a 100 × 1.46 NA alpha plan apochromat oil immersion objective and a pco.edge sCMOS camera. Fluorescence images were acquired sequentially using 200–300 ms exposure times per image, for a total of 15 images per SIM reconstruction. All imaging was performed at room temperature (∼23°C). Raw data were reconstructed using the SIM algorithms in ZEN 2011 SP7 software (black edition, Carl Zeiss). Brightfield images were captured using widefield imaging mode. Images had a final pixel size of 25 nm.

Gated STED (gSTED) images were acquired on a Leica TCS SP8 STED 3× system, using an HC PL Apo 100 × oil immersion objective with NA 1.40. Fluorophores were excited using a white excitation laser operated at 509 nm for mYFP and 563 nm for mKate2. A STED depletion laser line was operated at 592 nm and 775 nm for mYFP and mKate2, respectively, using a detection time-delay of 0.8–1.6 ns for both fluorophores. The total depletion laser intensity was in the order of 20–40 MW cm^−2^ for all STED imaging. The final pixel size was 13 nm and scanning speed was 400 Hz. The pinhole size was set to 0.9 AU.

Images were processed and analyzed using Fiji (71). Line scans were analyzed using the Plot Profile function in Fiji, using a line width of 1.5. Fluorescence intensities were normalized to the highest value for each channel.

### Scanning- and transmission electron microscopy analysis

Overnight cultures were diluted to approximately OD_600_ = 0.1. When OD_600_ reached 0.4, the cultures were diluted 1:250. Antibiotic and IPTG were added when appropriate. The cultures (10 ml) were grown to OD_600_ = 0.3 and 1 volume of fixation solution, containing 5 % (w/v) glutaraldehyde and 4 % (w/v) paraformaldehyde in 1×PBS, pH 7.4, was added. The tubes were carefully inverted a few times and incubated for one hour at room temperature before being placed at 4°C overnight. The following day, the cultures were centrifugated at 5000 × *g*, and the pellets washed three times with PBS. Further preparations of samples to be analyzed with TEM were performed as described before (69).

Samples for SEM were, after washing with PBS, dehydrated with EtOH, essentially in the same manner as for sample preparations for TEM (69). The samples were subjected to critical point drying by exchanging the EtOH with CO_2_. Then, the samples were coated with a conductive layer of Au-Pd before being analyzed in a ZEISS EVO50 EP Scanning electron microscope. Images were analyzed and prepared using Fiji (71).

### Growth assays

For examining growth on solid media, overnight cultures were diluted 1:250 in BHI containing 300 µM IPTG, unless otherwise specified. When reaching exponential phase, OD_600_were adjusted to 0.3 for all samples. A 10-fold dilution series were made for all strains, and 2 µl of each dilution were spotted on BHI agar containing proper antibiotics and 300 µM IPTG. The plates were incubated at 37°C for approximately 16 h, and pictures of the plates were captured in a Gel Doc™ XR + Imager (Bio-Rad).

For measurement of growth in liquid cultures, cells were at OD_600_ 0.4, were diluted 1:250 in medium containing 300 µM IPTG. Every hours for five hours, OD_600_ was measured spectrophotometrically using Genesys 30 (Thermo Scientific) and dilutions of the cultures were plated for CFU-counting.

### Minimum inhibitory concentration (MIC) assays

The experiments were set up in 96-well microtiter plates with a total volume of 300 µl. A two-fold dilution series of the antibiotics were prepared in BHI containing selective antibiotics and IPTG when appropriate. The overnight cultures were diluted 1:1000 in BHI containing 300 µg/ml IPTG for induction. The cells were grown at 37°C, and the plate was shaken for 5 sec before measurements of OD_600_ were taken every 10th min throughout the experiment, using either a Synergy™ H1 Hybrid Multi-Mode Reader (BioTek Instruments) or a Hidex Sense (Hidex Oy). The experiments were repeated at least two times with the same results.

### RNA isolation and RT-PCR

To verify that *smdA* expression was knocked down by CRISPRi, RNA was isolated from exponentially growing cultures of IM284 (SH1000, CRISPRi(control)), IM165 (SH000 CRISPRi(empty)) and IM269 (SH1000 CRISPRi(*smdA*)). Isolation of total RNA and cDNA synthesis were performed as previously described (69). A PCR reaction (30 cycles) was run with Phusion® High-Fidelity DNA polymerase (NEB). The primer pairs im126/im127 and im137/im138 were used to target the reference gene *pta* (74) and *smdA*, respectively.

### Detection of lipoteichoic acid (LTA) by Western blotting

Detection of LTA was performed by western blotting, and sample preparations were done according to descriptions found in Hesser et al (26). The samples were separated on a 4-20 % gradient Mini Protean TGX acrylamide gel (BioRad), and subsequently transferred to a polyvinylidene difluoride (PVDF) membrane by semi-dry electroblotting. The membrane was blocked for 1 h in 5 % (w/v) skim milk in PBST and placed overnight at 4°C. Next, the membrane was incubated for 1 h with α-LTA (Hycult) 1:4000 in PBST, washed three times (10 minutes each) with PBST before incubation for 1 h with α-Mouse IgG HRP Conjugate (Promega) secondary antibody (1:10 000 in PBST). The membrane was again washed three times and LTA bands were visualized by using SuperSignal™ West Pico PLUS Chemiluminescent substrate (Thermo Fisher Scientific) in an Azure Imager c400 (Azure Biosystems).

Same procedure as in (44) was carried out for possible detection of LTA released to the medium. The supernatants, after harvesting cells during sample preparations, were kept. Supernatant samples were centrifugated for 16 000 × *g* for 10 min, and 75 µl was mixed with 25 µl 4 × SDS-PAGE sample buffer. These samples were boiled for 30 min and applied on the 4-20 % Mini Protean TGX acrylamide gel. Thereafter, the same immunodetection procedure as described above was followed.

### Fourier-transform infrared spectroscopy (FTIR) analysis

Cultures of the strains IM313 (HG001, CRISPRi(control)), IM312 (HG001, CRISPRi(*smdA*)) and IM357 (HG001, CRISPRi(*tarO*)) were initially pre-grown to exponential phase, back diluted to OD_600_ = 0.05 and induced with 300 µM IPTG. The bacterial cells (1 ml) were harvested at OD_600_ = 0.4 by centrifugation at 5000 × *g*, 4°C, for 3 min. The pelleted cells were kept at -20°C prior to further processing. Pellets were resuspended in 40 µl 0.1 % (w/v) NaCl, and 10 µl of the suspensions were added to an IR-light-transparent silicon 384-well microplate (Bruker Optic, Germany), with three technical replicates for each sample. The plates were left to dry at room temperature for approximately 2 h. FTIR spectra were recorded in transmission mode using a high-throughput screening extension (HTS-XT) unit coupled to a Vertex 70 FTIR spectrometer (Bruker Optik GmbH, Leipzig, Germany). Spectra were recorded in the region 4000-500 cm^-1^, with a spectral resolution of 6 cm^-1^, a digital spacing of 1.928 cm^-1^, and an aperture of 5 mm. For each spectrum, 64 scans were averaged. The OPUS software (Bruker Optik GmbH, Leipzig,Germany) was used for data acquisition and instrument control. The obtained spectra were processed by taking second derivatives and extended multiplicative signal correction (EMSC) preprocessing in Unscrambler X version 11 (CAMO Analytics, Oslo, Norway). Results presented are averaged spectra from 3 biological replicates (each with 3 technical replicates) for the region with wavelengths between 1200 cm^-1^ and 800 cm^-1^.

### GFP-trap and liquid chromatography with tandem mass spectrometry (LC-MS/MS)

Cultures (*S. aureus* SH1000 wild-type, IM308 SH1000 *smdA-m(sf)gfp*, IM104 SH1000 pLOW-*smdA-m(sf)gfp* and IM164 SH1000 pLOW-*smdA-flag*) were pre-grown to exponential phase, back diluted to OD_600_ = 0.05 and induced if necessary. When reaching OD_600_ at 0.4, 80 ml of each culture were harvested by centrifugation at 4000 × *g*, 4°C for 3 min. Supernatants were decanted and pellets resuspended in cold TBS prior to transfer to 1.5 ml microcentrifuge tubes. Centrifugation was repeated for 1 min, and pelleted cells were stored at -80°C prior to further use.

For GFP-trap, cells were resuspended in 1 ml cold buffer containing 10 mM Tris pH 7.5, 150 mM NaCl, 1 mM PMSF, 6 µg/ml RNase and 6 µg/ml DNase. Suspensions were transferred to 2 ml lysing matrix B tubes (MP Biomedicals) containing 0.8 g ≤ 106 µm glass beads (Sigma-Aldrich) and subjected for mechanical lysis by agitation in a FastPrep-24™ (MP Biomedicals) for 3 × 30 sec at 6.5 m/s, with 1 min pause on ice between the runs. Tubes were centrifugated at 5000 × *g*, 4°C for 10 min, and supernatants transferred to new tubes. Concentrations were determined measuring Abs280 using NanoDrop™ 2000 (Thermo Fisher Scientific), where a small amount of the samples were added a final concentration of 1 % (w/v) SDS prior to measurements. GFP-Trap beads (25 µl per sample) (Chromotek) were washed three times with 500 µl ice cold Dilution/Wash buffer (10 mM Tris pH 7.5, 150 mM NaCl, 0.5 mM EDTA) and centrifugated at 2500 × *g*, 4°C for 5 min. Lysates were diluted in Dilution/Wash buffer to a final concentration of 1 mg in a total volume of 500 µl, before being transferred to GFP-Trap beads. Samples were placed in a Bio RS-24 Multi rotor (Biosan) at 4°C for 1 h. Then, samples were centrifugated at 2500 × *g*, 4°C for 5 min, supernatant removed, and beads washed three times with Dilution/Wash buffer. During last washing step, solutions were transferred to new tubes, and after centrifugation and removal of supernatant, beads were resuspended in 50 µl 5 % (w/v) SDS, 50 mM Tris pH 7.6. Tubes were incubated at 95°C for 5 min, and centrifugated at maximum speed for 30 sec. After standing at the bench a few minutes, 30-50 µl were transferred into new tubes. Samples were kept at -20°C and heated for 2 min at 95°C prior to the sample preparation method Suspension trapping (STrap), conducted as described by A. Zougman et al. (75).

The peptide samples were analyzed by coupling a nano UPLC (nanoElute, Bruker) to a trapped ion mobility spectrometry/quadrupole time of flight mass spectrometer (timsTOF Pro, Bruker). The peptides were separated by an Aurora Series 1.6 μm C18 reverse-phase 25 cm × 75 μm analytical column with nanoZero and CaptiveSpray Insert (IonOpticks, Australia). The flow rate was set to 400 nl/min and the peptides were separated using a gradient from 2 % to 95 % acetonitrile solution (in 0.1 % (v/v) formic acid) over 120 minutes. The timsTOF Pro was ran in positive ion data dependent acquisition PASEF mode, with a mass range at 100-1700 m/z. The acquired spectra were analyzed against a *S. aureus* NCTC8325 proteome database.

### Bacterial two-hybrid (BACTH) assays

Plasmid construction, and procedure for the BACTH assays, were conducted in a same manner as previously described (69), and primers used are listed in Table S3. Briefly, gene fusions of selected genes, to the T18 or T25 domains of adenylate cyclase form *Bordetella pertussis*, were made by restriction cutting and ligation in the plasmid vectors pKT25, pKNT25, pUT18 or pUT18C (Euromedex). *E. coli* XL1-Blue cells were used for transformation, and plasmids verified by sequencing before BACTH assays (76) were set up according to the manufacturer (Euromedex). Co-transformation of plasmids containing fusion-genes of opposite domains, that is T25 in one plasmid and T18 in the other, were done in *E. coli* BTH101 with 50 µg/ml kanamycin and 100 µg/ml ampicillin as selection markers. Five random colonies were picked, grown in liquid LB to visible growth, and spotted on LA plates containing 40 µg/ml X-gal and 0.5 mM IPTG, in addition to the selection markers. Plates were incubated dark at 30°C for 20-48 h before being inspected, and blue colonies are an indication of positive interaction between tested genes. Presented results are representative for at least six independent replicates.

## Acknowledgements

We would like to acknowledge the NMBU Imaging Center for help with electron microscopy, Maria Victoria Heggenhougen and Marita Torrissen Mårli (both NMBU) for access to unpublished strains and Henriette Olsen (NMBU) for help with FTIR. Mass spectrometry-based proteomic analyses were performed by The MS and Proteomics Core Facility, Norwegian University of Life Sciences (NMBU). This facility is a member of the National Network of Advanced Proteomics Infrastructure (NAPI), which is funded by the Research Council of Norway INFRASTRUKTUR-program (project number: 295910). We acknowledge the van Nieuwenhze group, Indiana University, for providing HADA. The work is supported by grants from the Research Council of Norway (project number 250976) and JPI-AMR (project number 296906). Work in the Structural Cell Biology Unit (OIST) is supported by OIST core subsidy. Ine S. Myrbråten acknowledge support from “Pasteurlegatet”.

## Legends to supplementary material

**Fig. S1. Multiple sequence alignment of SmdA from different staphylococcal species**. Protein sequences were aligned with Clustal Omega (78). The blue shaded residues are predicted to be extracellular, the transmembrane domain is shaded in grey, and the predicted NERD domain is shaded in yellow. *S. aureus* NCTC8325-4 is highlighted in bold, and residues that were mutated are marked in green. The accession numbers of the sequences are indicated, and the first five letter in the sequence tags indicate the genus or species corresponding the to the sequences (for example, Nosoc; *Nococomiicoccus*, Aliic; *Aliicoccus*, Salin; *Salinicoccus*, Jeotg; *Jeotgalicoccus*, Auric; *Auricoccus*, Abyss; *Abyssicoccus*, Mepid; *Macrococcus epidermidis*, Mcase; *Macrococcus caseolyticus*, Ssciu; *Staphylococcus sciuri*, Sinte; *Staphylococcus intermedius*).

**Fig. S2. Growth of SmdA knockdown strains in liquid cultures and verification of *smdA* silencing. (A)** Growth of SmdA^down^ in *S. aureus* SH1000 (IM269) compared to the CRISRPi-control strain with a non-targeting sgRNA (IM284). IPTG (300 µg/ml) was added to induce expression of the CRISPRi-system and CFU/ml and OD_600_ measured every hour for five hours. (B)Verification of *smdA* silencing by PCR with RT-PCR. cDNA was synthesized from RNA isolated from induced and un-induced cultures of SH1000 SmdA^down^ (IM269) and the CRISPRi control strains (IM284; non-targeting sgRNA and IM165; empty plasmid without sgRNA). Primers targeting either *smdA* or the housekeeping gene *pta*.

**Fig. S3. Transmission electron microscopy (TEM) of different *S. aureus* strains with SmdA knockdown**. SmdA^down^ and control cells analyzed with TEM in the *S. aureus* strains (**A**) NCTC8325-4 (IM311 and IM307), (**B**) HG001 (IM312 and IM313) and (**C**) COL (IM294 and IM295). Th e sizes of the scale bars are indicated in the images.

**Fig. S4. Scanning electron microscopy (SEM) of cells depleted of SmdA**. SEM micrographs of SmdA^down^ and control cells in *S. aureus* (**A**) HG001 (IM312 and IM313) and (**B**) COL (IM294 and IM295). All scale bars, 1 µm.

**Fig. S5. Overexpression of SmdA**. (**A**) Induced expression of an ectopic copy of *smdA* in the plasmid pLOW in *S. aureus* NCTC8325-4 (MK1866). Cells were labelled with the cell wall marker fluorescent vancomycin (VanFL). Scale bar, 2 µm. (**B**) Autolysis of *S. aureus* SmdA overexpression strain (MK1866) compared to plasmid control strain (MK1465) monitored in presence of 0.5 % Triton X-100. Results presented as % of initial OD_600_. Error bars represent standard error calculated from four technical replicates.

**Fig. S6. Analysis of teichoic acids**. (**A-B**) Lipoteichoic acid (LTA) detection by immunoblotting with α-LTA antibody was performed with (**A**) cell extract samples from SmdA^down^ and CRISPRi control cells from *S. aureus* SH1000 (IM269 and IM284), NCTC8325-4 (IM311 and IM307), HG001 (IM312 and IM313) and COL (IM294 and IM295), and (**B**) cell extract (CE)- and supernatant (SN) samples from SmdA^down^ and CRISPRi control cells from *S. aureus* SH1000 and NCTC8325-4 (IM269, IM284, IM311 and IM307, respectively). In (A), the mean intensities in the bands (background subtracted) were determined using Fiji (71). The mean intensities of the LTA bands in the *smdA* depletions relative to their controls are plotted. All control strains express a non-targeting sgRNA. (**C-D**) SmdA does not have major impact on the synthesis of wall teichoic acids. (**C**) TEM micrographs of *S. aureus* NCTC8325-4 control strain (IM307) compared to SmdA and TarO knockdown strains IM311 and IM358, respectively. The black arrows indicate presence of a high-density layer in the septum, which is missing in the TarO depletion strain (white arrow). (**D**) Fourier transform infrared spectroscopy (FTIR) of *S. aureus* HG001 control strain (IM313) and knockdown strain of SmdA and TarO (IM312 and IM357, respectively). The polysaccharide region of the spectrum is shown, and the indicated peaks represents α- and β-glycosidic bonds.

**Fig. S7**. Bacterial two-hybrid analysis demonstrating self-interaction between SmdA proteins. Blue bacterial spots indicate positive interactions and white spots indicate no interaction.

**Table S1. Proteins pulled down with SmdA-GFP**.

**Table S2. Strains used in this study**.

**Table S3. Primers used in this study**.

