## Supplemental material for "SmdA is a novel cell morphology determinant in *Staphylococcus aureus*"

**(Myrbråten et al.)**

```
Nosoc_WP_068130731     ----------------------------------MSTEILTEPLFIIVAALLLIVIISFL   26
Aliic_SEV88016         ----------------------------------MNTDIFLDPIFIAVAAVLLIVLTWFL   26
Salin_WP_020007446     ----------------------------------MNTEIFTNPIVIAILAVLLIVAVSFL   26
Jeotg_WP_026866661     ----------------------------------MNTEIFTNPIFIAIAVVLLIVVIWFL   26
Auric_AQL56528         ------------------------------MNINMNDFDMIHWIMFGLALLLLILLIFFI   30
Abyss_RPF57660         ------------------------------MNINMNDFDMIHWIMFGLALLLLILLVFFI   30
Mepid_RAK44338         --------------------------------MSVEVGSTLFYVLIGLGVLALLFFVLWL   28
Mcase_PKE19057         MQFYFLINLSKTMKLDMFFFIMKLLNIRKVRAMSLSNLQPMHYIVLGLAAALLLFIVLFI   60
Ssciu_WP_025905540     ----------------------------------MSQLEPIHYGLIAAIVIAILFIILFF   26
Sinte_PNZ51430         ----------------------------------MTQFGPMEIGLIAAIVVAALFFILFL   26
Schro_KDP13111         ----------------------------------MSQLDPIQIGLIAVSVLALLFLVLFL   26
Shyic_RTX86857         -------------------------------------MGPLEIGLVAAIVIAVIFLILFL   23
Saur_NCTC8325          ----------------------------------MDLSSPIVIGLIIAVVIALIFFVLFL   26
Ssimi_EHJ07687         ----------------------------------MDFSSPIVIGLIAAVVIALIFFVLYL   26
Swarn_PTI58951         ----------------------------------MSLSSPIGIGLIVAVVIAIIFFVLFL   26
Scap_TPX81490          ----------------------------------MSLSSPIVIGLIVAIIVAILFFILFL   26
Mabsc_SLD57873         ----------------------------------MDFSSPTVIGLIVAIPVAVLFFVLFL   26
Sepid_KAB2218069       ----------------------------------MDFSSPTVIGLIVAILVAVLFFILFL   26
Sarle_EJY95024         ----------------------------------MNSFGPIEIGLIVAVIVAVICLILFL   26
Ssapr_EHY92709         ----------------------------------MNSFGPIEIGLIVAIVVAVICLILFI   26
Scarn_RTX88654         ----------------------------------MNSFGPIEIGLIAAIVVAIICLILFL   26
Slugd_ADC87241         ----------------------------------MNSLGPMEIGLIVAVVIALICLILFF   26
                                                                      . .      :   : :

Nosoc_WP_068130731     IYYVKHRNEVEENERLYKKKEETLIESYVKNQEDERMAHKKEVSHLNEKYLEDTTLLNNK   86
Aliic_SEV88016         VYYFRSRNRVNKLTEEFDHEKQTLIEDYEATQEEDRLSHKKEVSGLNEKYNKDTEQLNKR   86
Salin_WP_020007446     VYFLKNRNRINTLTDEYSKEKEGLIEKYESNQEEERLNHKKEVSTLNEKYHTDTTLLNNK   86
Jeotg_WP_026866661     VYYFKHRNQVKVVENNHAKEKESLVQKYESEHEAERLEHKKELSNLNEKYYNDTTLLDNK   86
Auric_AQL56528         VKMIKANKQYKELQQARDDREKKLTSDYEKRIETERVDGKKKFSEQQSKYDAIVDDQSSQ   90
Abyss_RPF57660         VKMIKANKQYKELQQARDDREKKLTSDYEKRIETERVDGKKKFSEQQSKYDAIVDDQSSQ   90
Mepid_RAK44338         LALSSKKKSIARKEEEFRKAQKIRSDEYHDESEKSRLQYKKELAEQKETLQKTIDEKSSH   88
Mcase_PKE19057         YALSSKRKAIRAKEEALNKERSEMKSNYEESSEKSRLTFKKELAEQKDTYEAQLSTQNAQ  120
Ssciu_WP_025905540     VSLKQKQKSLNKIQEAHKKENETLKSEHKEKLDHERVENKKVLTKQEETHQEAISQKERE   86
Sinte_PNZ51430         VALKSKKKAKQTYATQYQTKQDKLTHEHQEELEKVRIDKKKAETRHKEEYETMVSSKNRE   86
Schro_KDP13111         FALRSKKKAKETYANQYQSRETKLNNEHKEALEKARIEKKKSDTRHKEEYDTMVSSKNRE   86
Shyic_RTX86857         TALNSKKKAQQQAEEQYEAKEKSLKDNYEDELEKERVEHKKTVTKQRADFDATVDSKDRE   83
Saur_NCTC8325          VALGSKKKVKRQTEEKYEQQEQNIKKSHEEALEKERIQNKKTITKQQEDYNHMVSTKDRE   86
Ssimi_EHJ07687         VALGSKKKVKRQTEEKYEQKEQNIKKTHEEQLEKERIENKKTITKQQEDYNEMVSTKDRE   86
Swarn_PTI58951         IALNSKKKIKQQTEEEYQQKEQSIKASHEEALEKERIENKKTVTKQKEDYEATVNSKERE   86
Scap_TPX81490          VANHSKKKIKNQTEAQYKEKEQHMKKSHEEALEKERVENKKAVTKQKEDFDATVSSKDRE   86
Mabsc_SLD57873         VANHSKKKVKNQTEAHYKEKEQHLKESHEEALEKERVENKKVVTKQQEDFDVTVSNKNRE   86
Sepid_KAB2218069       VANHSKKKVKNQTEAHYKEKEQHLKESHEEALEKERVKNKKVVTKQKEDFDVTVSNKNRE   86
Sarle_EJY95024         VTLKSKKKAQQKAEEHYQKKEKQLQDSYAEELEKERIENKKTVTKQKEDYDHTVNSKNRE   86
Ssapr_EHY92709         VALKSKKKAQEKVEAQYKSREQQLSDEHEEELEKERIENKKTVTKQKEEYTAAVNSKDRE   86
Scarn_RTX88654         VTLKSKNNIKQNTKEEYSLKEKQMLESHEEALEKERIENKKQVTRQKEDFDATISGKNRE   86
Slugd_ADC87241         VALRNNKKIKRQTVDEYKLKEKQMQESHDEALEKERIENKKTVTKQKENYEATVNSKERE   86
                             .:             .      :    :  *:  **  :  .          . .

Nosoc_WP_068130731     LSSIQQFTVDKGEYLTDLALLNFKNKLVTEERIRESDMYILSNIYLPSRNYTNTRKIDHL  146
Aliic_SEV88016         LRSVSQFTSDKGEYLTDLALKDLKNQLVKEDKIRDLDMHILSNIYLPSRNYTNTRKVDHL  146
Salin_WP_020007446     LSSLRQFTVDKGEYLTDLSLIQLKERLVRDEKIRETDMHILSNVYLPSRNYTNTRKIDHL  146
Jeotg_WP_026866661     LSSLHQFSVDKGEYLTDLALIQLKDKLVKDEKIRESDMIILSNVFLPSRNYTNTRKIDHL  146
Auric_AQL56528         ISSLKQFTYGKSQYLTDITLLSFRDKLIDQERIRPEDMHVLANVLIPSKNYKQTKQVDHV  150
Abyss_RPF57660         ISSLKQFTYGKSQYLTDITLLSFRDKLIDQERIRPEDMHVLANVLIPSKNYKQTKQVDHV  150
Mepid_RAK44338         IESLKMFSKDKGEYLTDLTLIQLKEQFIREERIRPEDMHVLANIYIPGKRVKSTDKLDHV  148
Mcase_PKE19057         IDSLKLFSRDKGEYLTDLTLINLKDNLVAQERIRPEDMHVLANIYIPGKRVKSTDRLDHV  180
Ssciu_WP_025905540     IDSLKLFSKNEGEYITDRHLLELRDQLVNERRIRPEDMHIMANIFLPKDPLGKVRQIDHL  146
Sinte_PNZ51430         IDALKLFSKNHSEYVTDMRLLGIRERLVKEKRIRPEDMHIMANIFLPTNDLEDITRVSHL  146
Schro_KDP13111         IDALKLFSKNDSEYITDMRLLGIRERLVKEKRIRPEDMHIMANIFMPTNDLEEITRISHL  146
Shyic_RTX86857         IDALKLFSKNHSEYITDMRLIGIRERLVKEKRIRPEDMHIMANIFLPKNDMNDIERISHL  143
Saur_NCTC8325          IDALKLFSKNHSEYVTDMRLIGIRERLVKEKRIRPEDMHIMANIFLPKDGFNNIERISHL  146
Ssimi_EHJ07687         IDALKLFSKNHSEYVTDMRLIGIRERLVKEKRIRPEDMHIMANIFLPSNKFNDIERISHL  146
Swarn_PTI58951         IDALKLFSKNHSEYVTDLRLIGIRERLVNEKRIRPEDMHIMANIFLPTNELNKIERISHL  146
Scap_TPX81490          IDALKLFSKNHSEYVTDMRLIGIRERLVNEKRIRPEDMHIMANIFLPSNEFNNIERISHL  146
Mabsc_SLD57873         IDALKLFSKNHSEYVTDMRLIGIRERLVNEKRIRPEDMHIMANIFLPSNELTNIERVSHL  146
Sepid_KAB2218069       IDALKLFSKNHSEYVTDMRLIGIRERLVNEKRIRPEDMHIMANIFLPSNELTNIERVSHL  146
Sarle_EJY95024         IDALKLFSKNHSEYVTDMRLIGIRERLVNEKRIRPEDMHIMANIFLPRNEFSDVQRISHL  146
Ssapr_EHY92709         IDALKLFSKNQSEYVTDMRLIGIRERLVNEKRIRPEDMHIMANIFLPRNEFSDVQRISHL  146
Scarn_RTX88654         IDALKLFSKNTSEYVTDMRLIGIRDRLVNEKRIRRDDMHIMANIFLPSNEFNDIQRISHL  146
Slugd_ADC87241         IDALKLFSKNTSEYVTDMRLIGIRERLVNEKRIRPEDMHIMANIFLPGNDLNNIERISHL  146
                       : ::  *: . .:*:**  *  :::.:: : :**  ** :::*: :*     .  ::.*:
```

```
Nosoc_WP_068130731      VLTRTGIYLIDSKYWSGHILHGVTEEQYDEIPYLEGIFQLLNLDPNKEQTLIFEKEND--    204
Aliic_SEV88016          VLTRTGIYLMESRFWKGHIIHGVSEQNFEQLPYVENFFELLGLNKKQEQTFIFEKKDD--    204
Salin_WP_020007446      VLTRTGIYMIDSKYWRGHILHGINEENFEELPYTESFFDLLELDKTKEQTLIFEKSDS--    204
Jeotg_WP_026866661      VLTRTGIYLIDSKYWSGHILHGVNEAQFETVPYVESFFDLLDLDKREQTLIFEKADQ--    204
Auric_AQL56528          ILTRTGIYIVDSNYFSGHVYHGMNEQQFDQFPFLEGVYDALGYDHKDEYSFIVEPKDN--    208
Abyss_RPF57660          ILTRTGIYIVDSNYFSGHVYHGMNEQQFDQFPFLEGVYDALGYDHKDEYSFIVEPKDN--    208
Mepid_RAK44338          VLTRTGIYLIDSNYWTGHIYHGVSEMQFAGEPMLEGVFNILELDPKLEQTIVLDKNKD--    206
Mcase_PKE19057          ILTRTGIYVIDSNYWTGHLYHGISEMQFDGEPIFETVFNILGLDPKSEQTICLDKAED--    238
Ssciu_WP_025905540      VLTRTGIYVIDSNLVSGHIYHGITEQQFSDFPVLGQVFETLDLNPNKEQTLLLEKQPN-N    205
Sinte_PNZ51430          VLTRTGLYVIDSELLKGHVYQGISQQQFADNPMMEQVFKTLNLSPQTPQTVVLDQSEAAQ    206
Schro_KDP13111          VLTRTGLYIIDSELLKGHVYQGVSQNQFRENPMMEHVFKTLHLDGQTPQTIVLDQKED-Q    205
Shyic_RTX86857          VLTRTGLYIIDSQLLKGHVYNGISGKQFAELPTIEQVFNVLNLDQRTPQTLVLDENDD-Q    202
Saur_NCTC8325           VLTRTGLYIIDSQLLKGHVYNGISGGQFKDLPPMEQVFDTLDLDKSRPQTIVMDQNDD-K    205
Ssimi_EHJ07687          VLTRTGLYIIDSQLLKGHVYNGISGGQFKDLPPMEQVFKTLDLDASRPQTIIMDQNDD-Q    205
Swarn_PTI58951          VLTRTGLYIIDSQLLKGHVYNGISGAQFSELPTMEQVFNTLELDKGTPQTLVLDQNSD-E    205
Scap_TPX81490           VLTRTGLYIIDSQLLKGHVYNGISGAQFKELPTMSQVFQTLDLDESTPQTLVLDQNED-Q    205
Mabsc_SLD57873          VLTRTGLYIIDSQLLKGHVYNGISGAQFKELPTMSQVFDTLDLDSSQPQTLVLDQNED-Q    205
Sepid_KAB2218069        VLTRTGLYIIDSQLLKGHVYNGISGAQFKELPTMSQVFDTLDLDSSQPQTLVLDQNED-Q    205
Sarle_EJY95024          VLTRTGLYIIDSQLLKGHVYNGISGAQFEETPMMSQVFDTLDLDKKDPQVLVLDQNDD-T    205
Ssapr_EHY92709          VLTRTGLYIIDSQLLKGHVYNGVSAAQFKEQPMMEQVFNTLDLDGQVPQTLVLDQNED-K    205
Scarn_RTX88654          VLTRTGLYIIDSQLLKGHVYNGISGNQFQELPMMEQVFNTLDLDPKSPQTLVLDQNDD-K    205
Slugd_ADC87241          VLTRTGLYIIDSKLLKGHVYNGISGNQFNELPMMQQVFSTLDLNAKAPHTLVLDQNDD-Q    205
                        :*****:*:::*.   **: :*:.  ::   *    .:. *  .      . .:

Nosoc_WP_068130731      EKVAVNYYNNTIEDTEMTAEKLRNVLKLQFDVVPIIYFNPKDNGGHTIMNYATNSNAKVI    264
Aliic_SEV88016          LNVTVSHFNHMIEEAKLTAEKLKNVLKLQFDVVPVVFFNPEDNGNYSIANYSNDPSVKVL    264
Salin_WP_020007446      KNVSVNHYNDTIDETKTTAEKLRNVLKLEYDIVPMIYFNPKDNGNYSITNYSADPSVKVL    264
Jeotg_WP_026866661      KNVSVNHYNGVIDETKITAEKLKNKLKLQYDIVPIIYFNPKDNGNYSITNYSEDPSIRVI    264
Auric_AQL56528          GEVVMHPLDHQIQDLKVTAEKIRNILKLQYPVKTIMFYNEQETKRNVSINYSTDKDVIVL    268
Abyss_RPF57660          GEVVMHPLDHQIQDLKVTAEKIRNILKLQYPVKTIMFYNEQETKRNVSINYSTDKDVIVL    268
Mepid_RAK44338          GNAEFRTYNHSIEALKLKAERLEKVFELPYPVTPIAYYNAEGVGSEHITNYSRDNGVKVI    266
Mcase_PKE19057          GSAQFKSYKHSIEELKLKAERLEKVFELPYPVTPIAYYNAEGVGSEHITNYSRDNSVKVI    298
Ssciu_WP_025905540      KTAAFHSYTDEVKEVDNTTEELQRQLELKYTPTPIIYFHPNEVSEATISNYSQDPNIKVL    265
Sinte_PNZ51430          NTLSVVDYTDRLAHVEALAVKLQKELDLKYTPTPILYFNPRHEGDVTISNYASHGVTKVL    266
Schro_KDP13111          QKLTVIDYTKHLDSVEKLATTLQQKLNLKYAPTPILYFNPRHEGDITISNYAPQSGTKVL    265
Shyic_RTX86857          QSATFVNYSKQLHYVEQLADALQRQLNLKYTPMALLYFNPKNEGAVTISNYAQNTNTKVL    262
Saur_NCTC8325           RSLSFVNYSDQIEAIKQLAEDLQKELGAKYTPTSILYFNLKNEGDVTISNYNQNSAVKVL    265
Ssimi_EHJ07687          KSLSFVNYSDKLESIKQLAEDLQKELGTKYTPTSILYFNPKNDGDVTISNYNQNAAVKVL    265
Swarn_PTI58951          KSLSFINYSDEIQQIQNVAGDLQSKLNTKYTPTSILYFNPKNEGDVTISNYAQNSNVKVL    265
Scap_TPX81490           QSLSFVNYSEKIKHIQNLAGDLQSELNTKYTPTSILYFNPKNDGDVTISNYNQASNVKVL    265
Mabsc_SLD57873          HSLSFVNYSDKIKHIEKLAGDLQNELNTKYTPTSILYFNPKKDNDVTISHYTQSSNVKVL    265
Sepid_KAB2218069        HSLSFVNYSDKIKHIEKLAGDLQNELNTKYTPTSILYFNPKKDNDVTISHYTQSSNVKVL    265
Sarle_EJY95024          QSLSVVNYSNHLEAVEKLAGDLQNELNLKYTPTTILYFNPKNDGGVTISNFAQTANSKVL    265
Ssapr_EHY92709          DALSFVNYTTHLNEIERLAGDIQTELNLKFTPTTILYFNPKNDGDVTITNYAQSSNSKVL    265
Scarn_RTX88654          SSLSFVNYSDQLEAIEKLATDLQTQLGTKYTPTSIMYFNPKHDGDVTISNYAQSSSVKVL    265
Slugd_ADC87241          SSLSFLDYSAQLAQIEKLATDLQTKLGLTYTPTSILYFNPKREGEVTISNYAQTSSAKVL    265
                            .      :   .  :  :.  :   :    : ::: .        ::       *:

Nosoc_WP_068130731      VGTEQLEDYFNKHVFQGRFQYTVKDLDEIAESLMRLNP    302
Aliic_SEV88016          IGNEELDEYFKKYVFHGRFQYTVKDLTDIVDQLKALNP    302
Salin_WP_020007446      VGEEELEAFFLKYVFHGRFQYTVKQLDEIAEEIFNLNP    302
Jeotg_WP_026866661      VGQEDLEAFFLKYVFHGRFQYTVKDLDEIADAILEQSL    302
Auric_AQL56528          MGKPELEEYFEKHVFHGRFEYTVEELEQIKQQLLEMNP    306
Abyss_RPF57660          MGKPELEEYFEKHVFHGRFEYTVEELEQIKQQLLEMNP    306
Mepid_RAK44338          VGEEELQTFFEKFVFHGRFQYKVEALENIMDQIEHLNP    304
Mcase_PKE19057          VGEKELEHFFEKFVFHGRFQYKVEELEEVRAAIEHLNP    336
Ssciu_WP_025905540      VGEKQLQHFFNKFVFHGRFQYSVEDLERIMDEIEKFNP    303
Sinte_PNZ51430          VGPEQLNEFFNRFVFHGRIQYDVNALQNIMDEIESFN-    303
Schro_KDP13111          VGPAQLDEFFNKFVFHGRIQYDVNDLQAIMDEIEAFN-    302
Shyic_RTX86857          VGAEQLDEYFNKFVFHGRIQYNVEDLARIMEEIESFN-    299
Saur_NCTC8325           VGAEQLDEFFNKFVFHGRIQYNVEDLQQMMDKIESFN-    302
Ssimi_EHJ07687          VGKEQLDEFFNKFVFHGRIQYNVDDLQNIMDKIESFN-    302
Swarn_PTI58951          VGPEQLDEFFNKFVFHGRIQYNVDDLQQIMEQIESFN-    302
Scap_TPX81490           VGPEQLDEFFNKFVFHGRIQYNVDELQTIMDKIESFN-    302
Mabsc_SLD57873          VGPEQLEEFFNKFVFHGRIQYNVDDLQDIMDKIESFN-    302
Sepid_KAB2218069        VGPEQLDEFFNKFVFHGRIQYNVDDLQDIMDKIESFN-    302
Sarle_EJY95024          VGPEQLDEYFNKFVFHGRIQYNVEDLQQVMDKIESFN-    302
Ssapr_EHY92709          VGPEQLDEFFNKFVFHGRIQYNVEDLQRVMDEIESFN-    302
Scarn_RTX88654          VGPEQLDEFFNKFVFHGRIQFNVEELQSIMDKIETFN-    302
Slugd_ADC87241          VGPEQLNEFFNKFVFHGRIQYNVDDLQEIMTKIESFN-    302
                        :*  :*: :* :.**:**::: *. *  :   :   .
```

**Fig. S1. Multiple sequence alignment of SmdA from different staphylococcal species.** Protein sequences were aligned with Clustal Omega (1). The blue shaded residues are predicted to be extracellular, the transmembrane domain is shaded in grey, and the predicted NERD domain is shaded in yellow. *S. aureus* NCTC8325-4 is highlighted in bold, and residues that were mutated are marked in green. The accession numbers of the sequences are indicated, and the first five letter in the sequence tags indicate the genus or species corresponding the to the sequences (for example, Nosoc; *Nococomiicoccus*, Aliic; *Aliicoccus*, Salin; *Salinicoccus*, Jeotg; *Jeotgalicoccus*, Auric; *Auricoccus*, Abyss; *Abyssicoccus*, Mepid; *Macrococcus epidermidis*, Mcase; *Macrococcus caseolyticus*, Ssciu; *Staphylococcus sciuri*, Sinte; *Staphylococcus intermedius*).

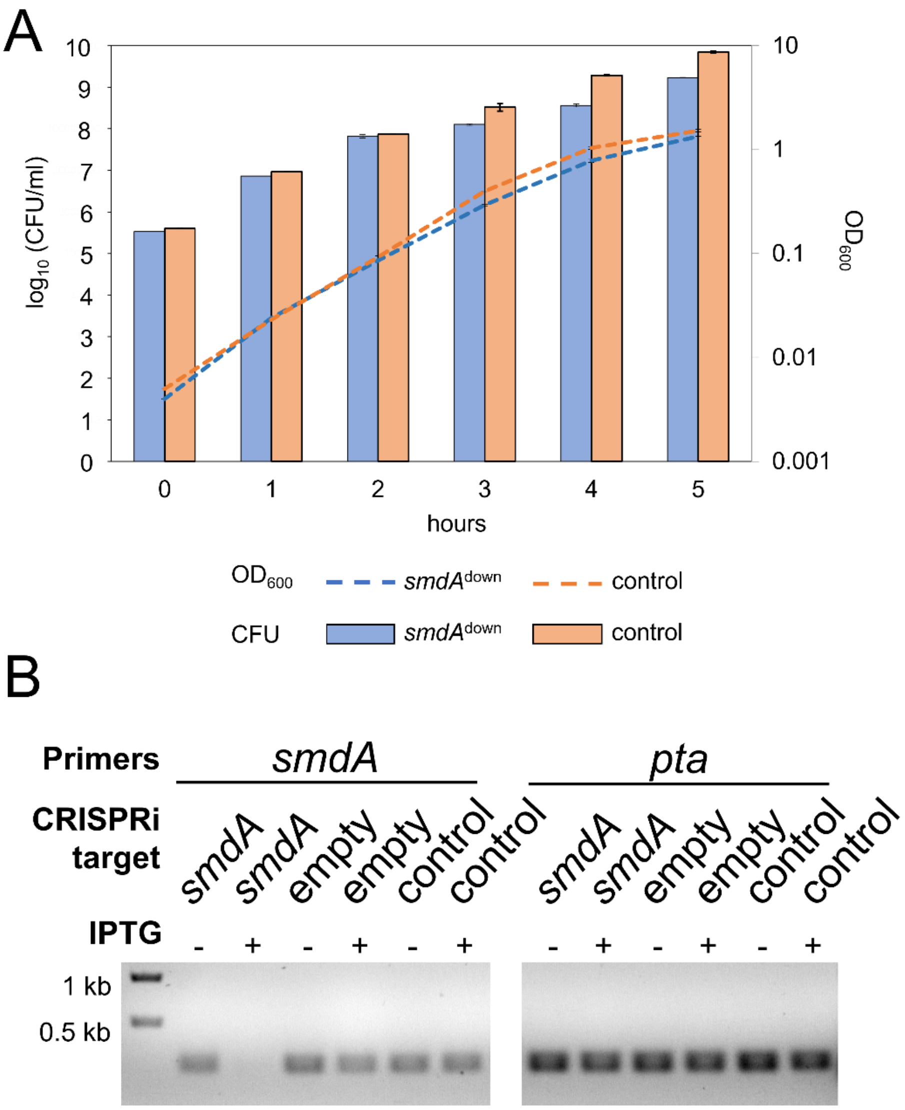

**Fig. S2. Growth of SmdA knockdown strains in liquid cultures and verification of *smdA* silencing. (A)** Growth of SmdA$^{down}$ in *S. aureus* SH1000 (IM269) compared to the CRISRPi-control strain with a non-targeting sgRNA (IM284). IPTG (300 µg/ml) was added to induce expression of the CRISPRi-system and CFU/ml and $OD_{600}$ measured every hour for five hours. (**B**) Verification of *smdA* silencing by PCR with RT-PCR. cDNA was synthesized from RNA isolated from induced and un-induced cultures of SH1000 SmdA$^{down}$ (IM269) and the CRISPRi control strains (IM284; non-targeting sgRNA and IM165; empty plasmid without sgRNA). Primers targeting either *smdA* or the housekeeping gene *pta*.

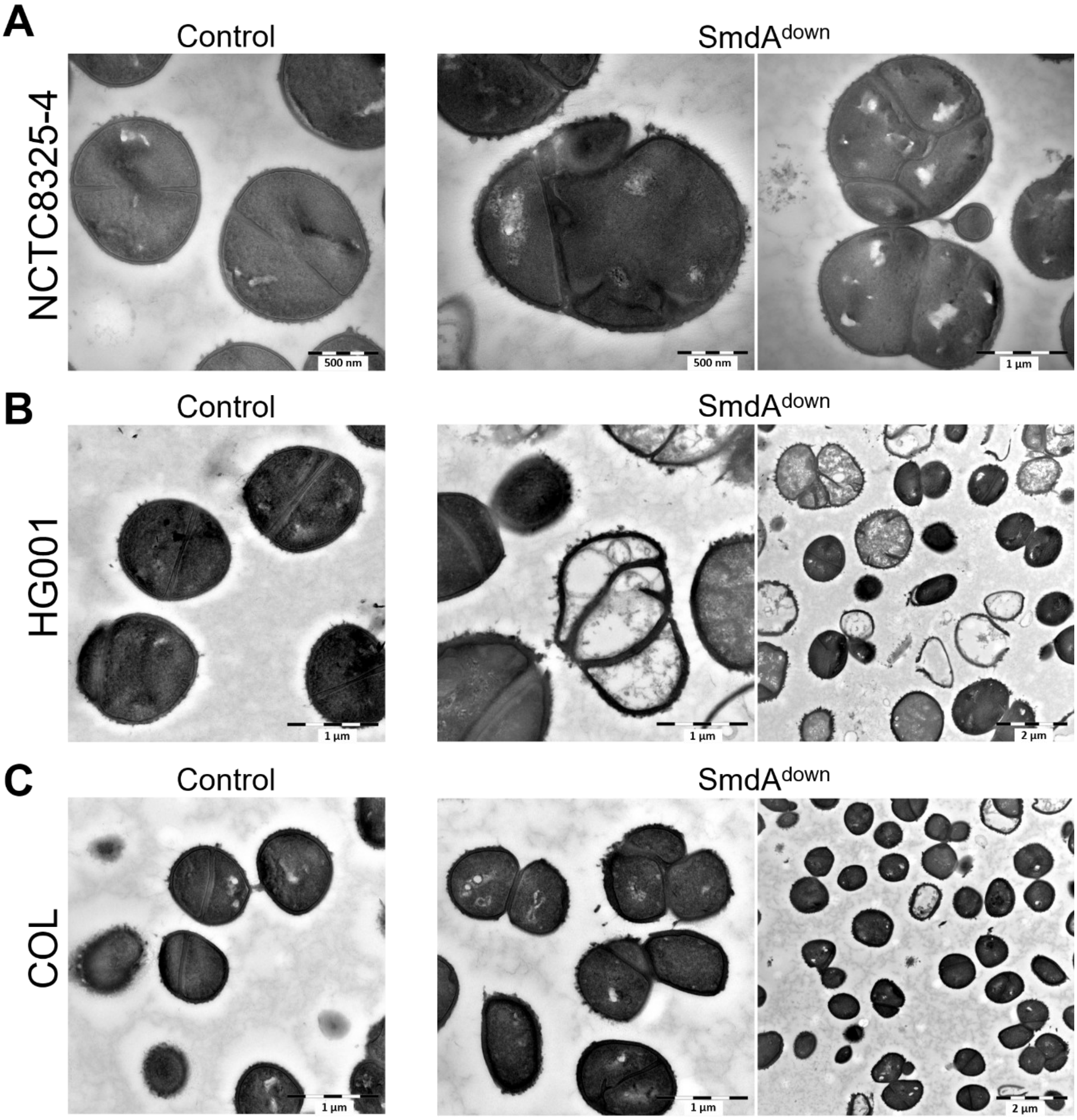

**Fig. S3. Transmission electron microscopy (TEM) of different *S. aureus* strains with SmdA knockdown.** SmdA$^{down}$ and control cells analyzed with TEM in the *S. aureus* strains (**A**) NCTC8325-4 (IM311 and IM307), (**B**) HG001 (IM312 and IM313) and (**C**) COL (IM294 and IM295). The sizes of the scale bars are indicated in the images.

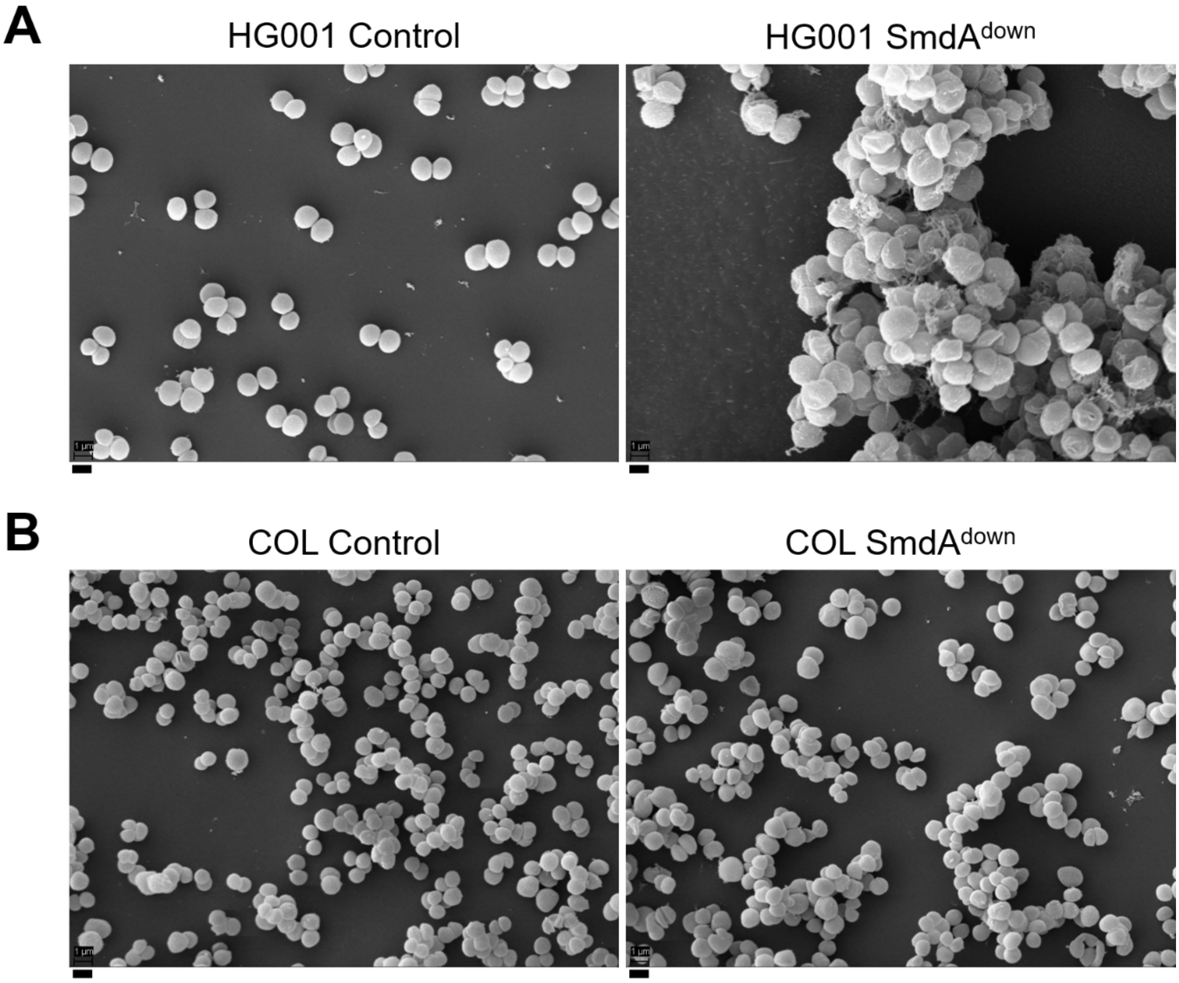

**Fig. S4. Scanning electron microscopy (SEM) of cells depleted of SmdA.** SEM micrographs of SmdA$^{down}$ and control cells in *S. aureus* (**A**) HG001 (IM312 and IM313) and (**B**) COL (IM294 and IM295). All scale bars, 1 µm.

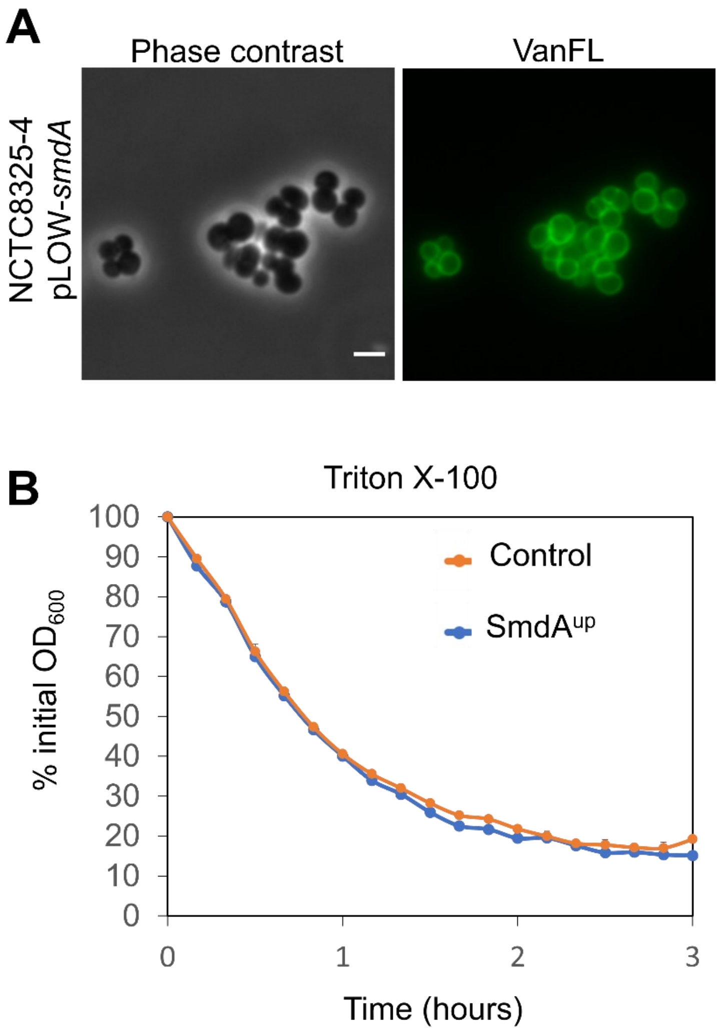

**Fig. S5. Overexpression of SmdA.** (**A**) Induced expression of an ectopic copy of *smdA* in the plasmid pLOW in *S. aureus* NCTC8325-4 (MK1866). Cells were labelled with the cell wall marker fluorescent vancomycin (VanFL). Scale bar, 2 µm. (**B**) Autolysis of *S. aureus* SmdA overexpression strain (MK1866) compared to plasmid control strain (MK1465) monitored in presence of 0.5 % Triton X-100. Results presented as % of initial $OD_{600}$. Error bars represent standard error calculated from four technical replicates.

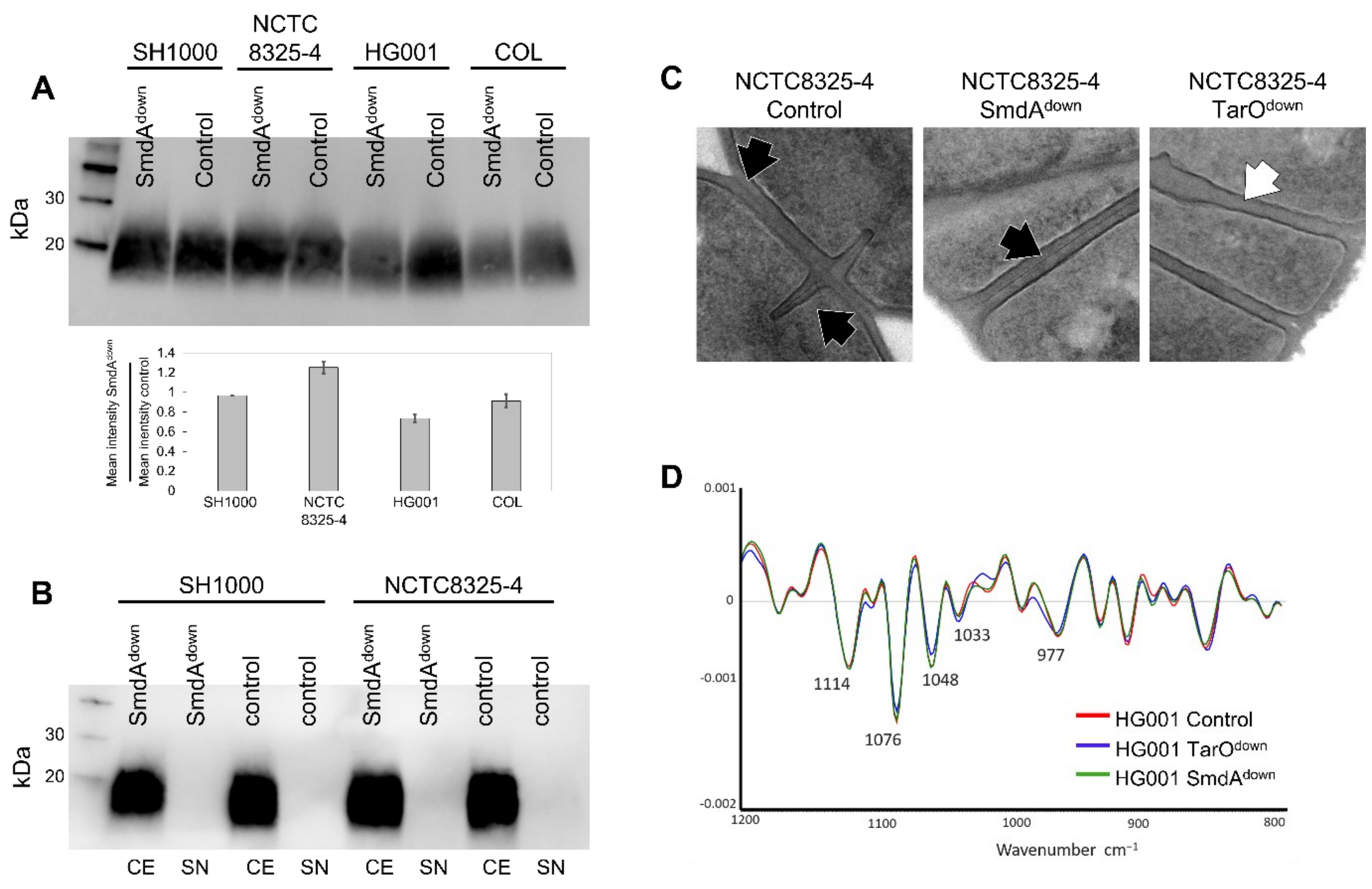
A
SH1000
NCTC 8325-4
HG001
COL
SmdAdown
Control
kDa
30
20
Mean intensity SmdAdown / Mean inentsity control
1.4
1.2
1
0.8
0.6
0.4
0.2
0
SH1000
NCTC 8325-4
HG001
COL
B
SH1000
NCTC8325-4
SmdAdown
control
CE
SN
C
NCTC8325-4 Control
NCTC8325-4 SmdAdown
NCTC8325-4 TarOdown
D
0.001
0
-0.001
-0.002
1114
1076
1048
1033
977
HG001 Control
HG001 TarOdown
HG001 SmdAdown
1200
1100
1000
900
800
Wavenumber cm-1

**Fig. S6. Analysis of teichoic acids.** (**A-B**) Lipoteichoic acid (LTA) detection by immunoblotting with α-LTA antibody was performed with (**A**) cell extract samples from SmdA$^{down}$ and CRISPRi control cells from *S. aureus* SH1000 (IM269 and IM284), NCTC8325-4 (IM311 and IM307), HG001 (IM312 and IM313) and COL (IM294 and IM295), and (**B**) cell extract (CE)- and supernatant (SN) samples from SmdA$^{down}$ and CRISPRi control cells from *S. aureus* SH1000 and NCTC8325-4 (IM269, IM284, IM311 and IM307, respectively). In (A), the mean intensities in the bands (background subtracted) were determined using Fiji (2). The mean intensities of the LTA bands in the *smdA* depletions relative to their controls are plotted. All control strains express a non-targeting sgRNA. (**C-D**) SmdA does not have major impact on the synthesis of wall teichoic acids. (**C**) TEM micrographs of *S. aureus* NCTC8325-4 control strain (IM307) compared to SmdA and TarO knockdown strains IM311 and IM358, respectively. The black arrows indicate presence of a high-density layer in the septum, which is missing in the TarO depletion strain (white arrow). (**D**) Fourier transform infrared spectroscopy (FTIR) of *S. aureus* HG001 control strain (IM313) and knockdown strain of SmdA and TarO (IM312 and IM357, respectively). The polysaccharide region of the spectrum is shown, and the indicated peaks represents α- and β-glycosidic bonds.

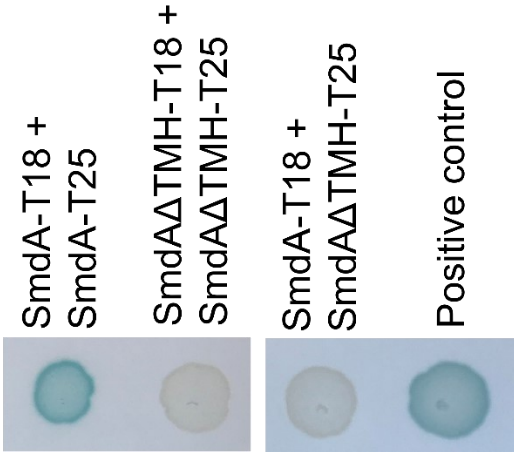

**Fig. S7. Bacterial two-hybrid analysis demonstrating self-interaction between SmdA proteins.** Blue bacterial spots and plus symbols indicate positive interactions and white spots and minus symbols indicate no interaction.

### Supplementary Tables

**Table S1. Proteins pulled down with SmdA-GFP.**

| Protein | Description | Chromosomal *smdA-gfp*[a] | | Plasmid-based *smdA-gfp* | |
|---|---|---|---|---|---|
| | | Unique peptides | Ratio | Unique peptides | Ratio |
| **SmdA** | Uncharacterized protein | 4 | >4 | 35 | 8.8 |
| **Atl** | Bifunctional autolysin | 4 | 4 | 29 | 5.8 |
| **SAOUHSC_01193** | Uncharacterized protein | 3 | >3 | 10 | 2.5 |
| **FtsH** | ATP-dependent zinc metalloprotease | 2 | >2 | 22 | 11 |
| **PrsA** | Foldase protein | 2 | >2 | 14 | 7 |
| **SpsB** | Signal peptidase I | 2 | >2 | 12 | >12 |
| **FhuD2** | Heme ABC transporter | 2 | >2 | 12 | >12 |
| **Pbp2** | Penicillin-binding protein 2 | 1 | >1 | 23 | 23 |
| **SAOUHSC_01676** | UPF0365 protein | 1 | >1 | 14 | 14 |
| **SAOUHSC_00356** | Uncharacterized protein | 1 | >1 | 14 | >4 |
| **FruA** | Fructose specific permease, putative | 1 | >1 | 12 | >12 |
| **AtpA** | ATP synthase subunit alpha | 1 | >1 | 12 | >12 |
| **AlaS** | Alanine--tRNA ligase | 1 | >1 | 12 | 12 |
| **GlpK** | Glycerol kinase | 1 | >1 | 12 | 12 |
| **NrdE** | Ribonucleoside-diphosphate reductase | 1 | >1 | 10 | >10 |
| RpoC | DNA-directed RNA polymerase subunit beta' | ND | NA | 28 | 9.3 |
| RpoB | DNA-directed RNA polymerase subunit beta | ND | NA | 24 | 12 |
| SAOUHSC_02525 | Uncharacterized protein | ND | NA | 18 | >18 |
| Pyk | Pyruvate kinase | ND | NA | 17 | 5.7 |
| AtpD | ATP synthase subunit beta | ND | NA | 15 | 7.5 |
| EzrA | Septation ring formation regulator | ND | NA | 15 | >15 |
| GlpD | Aerobic glycerol-3-phosphate dehydrogenase | ND | NA | 14 | 14 |
| SAOUHSC_01895 | Uncharacterized protein | ND | NA | 14 | >14 |
| Pbp1 | Penicillin-binding protein 1 | ND | NA | 14 | >14 |
| Pbp3 | Penicillin-binding protein 3 | ND | NA | 14 | >14 |
| ClpB | Chaperone protein | ND | NA | 14 | >14 |
| Mqo | Probable malate:quinone oxidoreductase | ND | NA | 13 | >13 |
| SecD | Multifunctional fusion protein | ND | NA | 13 | >13 |
| SAOUHSC_01974 | Uncharacterized protein | ND | NA | 13 | >13 |
| DltD | Protein DltD | ND | NA | 12 | >12 |
| ClpC | ATP-dependent Clp protease ATP-binding subunit | ND | NA | 12 | >12 |
| QoxA | Probable quinol oxidase subunit 2 | ND | NA | 11 | 11 |
| GatB | Aspartyl/glutamyl-tRNA(Asn/Gln) amidotransferase | ND | NA | 11 | >11 |
| SAOUHSC_00253 | Uncharacterized protein | ND | NA | 11 | >11 |
| SAOUHSC_00749 | Uncharacterized protein | ND | NA | 10 | >10 |
| Rny | Ribonuclease Y | ND | NA | 10 | >10 |
| ArgG | Argininosuccinate synthase | ND | NA | 10 | >10 |
| AccC | Biotin carboxylase | ND | NA | 10 | >10 |
| HtrA1 | Serine protease | ND | NA | 10 | >10 |

[a] ND; not detected, NA; not applicable.

**Table S2. Strains used in this study.**

| Name | Genotype[a] | Source |
|---|---|---|
| ***S. aureus*** | | |
| NCTC8325-4 | MSSA strain, derivative of NCTC8325, cured of phages. | (3) |
| SH1000 | *rbsU*+ derivative of strain NCTC8325-4 | (4) |
| RN4220 | Restriction deficient derivative of NCTC8325-4 | (5) |
| HG001 | MSSA-strain, derivative of NCTC8325 | (6) |
| COL | Hospital-associated MRSA strain | (7) |
| **CRISPRi depletion strains** | | |
| SAMK13 | SH1000, pLOW-*dcas9* | (8) |
| IM269 | SAMK13, pCG248-sgRNA*(smdA*), ery$^r$, cam$^r$ | This study |
| IM165 | SH1000, pLOW-*dcas9*, pCG248(empty), ery$^r$, cam$^r$ | This study |
| SAMK15/IM284 | SAMK13, pCG248-sgRNA*(*control), ery$^r$, cam$^r$ | (8) |
| IM307 | NCTC8325-4, pLOW-*dcas9*_extra_*lacO*, pCG248-sgRNA(control), ery$^r$, cam$^r$ | This study |
| IM311 | NCTC8325-4, pLOW-*dcas9*, pCG248-sgRNA*(smdA*), ery$^r$, cam$^r$ | This study |
| IM313 | HG001, pLOW-*dcas9*, pCG248-sgRNA*(*control), ery$^r$, cam$^r$ | This study |
| IM312 | HG001, pLOW-*dcas9*, pCG248-sgRNA*(smdA*), ery$^r$, cam$^r$ | This study |
| IM294 | COL, pLOW-*dcas9_aad9*, pCG248-sgRNA*(smdA*), spc$^r$, cam$^r$ | This study |
| IM295 | COL, pLOW-*dcas9_aad9*, pCG248-sgRNA*(*control), spc$^r$, cam$^r$ | This study |
| IM358 | NCTC8325-4, pLOW-*dcas9*, pCG248-sgRNA*(tarO*), ery$^r$, cam$^r$ | This study |
| IM357 | HG001, pLOW-*dcas9*, pCG248-sgRNA*(tarO*), ery$^r$, cam$^r$ | This study |
| IM293 | SH1000, pLOW-*dcas9*_P*atl-luc*, pCG248-sgRNA(*walR*) , ery$^r$, cam$^r$ | This study |
| **Strains for localization studies** | | |
| IM104 | SH1000, pLOW-*SAOUHSC_01908-m(sf)gfp,* ery$^r$ | This study |
| IM305 | NCTC8325-4, pLOW-*smdA-m(sf)gfp,* ery$^r$ | This study |
| IM373 | NCTC8325-4, pLOW-*smdAΔTMH-m(sf)gfp,* ery$^r$ | This study |
| IM308 | SH1000, *smdA-m(sf)gfp_aad9,* spc$^r$ | This study |
| HC060 | SH1000, pLOW-*smdA-mYFP*, pHC-*ftsZ-mKate2,* ery$^r$, neo$^r$ | This study |
| SH4639 | SH1000, *ezrA-gfp*, kan$^r$ | (9) |
| MK1952 | SH4639, pLOW-*dcas9*, pCG248-sgRNA(*smdA*), ery$^r$, cam$^r$ | This study |
| MK1953 | SH4639, pLOW-*dcas9*, pCG248-sgRNA(control), ery$^r$, cam$^r$ | This study |
| **Strains used for overexpression and mutagenesis** | | |
| MK1465 | NCTC8325-4, pLOW-dcas9, ery$^r$ | This study |
| MK1866 | NCTC8325-4, pLOW-*smdA*, ery$^r$ | This study |
| MK1911 | NCTC8325-4, pLOW-*smdAΔTMH*, ery$^r$ | This study |
| IM377 | NCTC8325-4, pLOW-*smdAΔTMH_mut1* (H145A), ery$^r$ | This study |
| IM378 | NCTC8325-4, pLOW-*smdAΔTMH_mut2* (R150A, T151A), ery$^r$ | This study |
| IM379 | NCTC8325-4, pLOW-*smdAΔTMH_mut3* (F280A, H281A), ery$^r$ | This study |
| **Other strains** | | |
| IM164 | SH1000, pLOW-*smdA-flag*, ery$^r$ | Lab collection |

| | | |
|---|---|---|
| ***E. coli*** | | |
| IM08B | DH10B, Δ*dcm*, $P_{help}$-*hsdMS*, $P_{N25}$-*hsdS* (expressing the *S. aureus* CC8 specific methylation genes) | (10) |
| BTH101 | Used for BACTH analysis | Euromedex |
| XL1-Blue | Host strain | Agilent |
| **Strains harboring plasmids used to facilitate cloning** | | |
| IM6 | IM08B, pLOW-*ftsZ-m(sf)gfp*, $amp^r$ | This study |
| IM98 | IM08B, pLOW-*ftsZ-m(sf)gfp*_KpnI, $amp^r$ | This study |
| IM33 | IM08B, pLOW-*lacA-m(sf)gfp*, $amp^r$ | This study |
| IM7 | IM08B, pLOW-*ftsZ-mYFP*, $amp^r$ | This study |
| IM8 | IM08B, pLOW-*ftsZ-mKate2*, $amp^r$ | This study |
| IM187 | IM08B, pMAD- *smdA-flag_aad9*, $amp^r$ | This study |
| **Strains used for BACTH assays** | | |
| GS1225 | XL1-Blue, pKNT25-*smdA*, $kan^r$ | This study |
| GS1226 | XL1-Blue, pUT18-*smdA*, $amp^r$ | This study |
| GS1302 | XL1-Blue, pKNT25-*smdAΔTMH*, $kan^r$ | This study |
| GS1303 | XL1-Blue, pUT18- *smdAΔTMH*, $amp^r$ | This study |
| GS1134 | XL1-Blue, pKT25-*pbp1*, $kan^r$ | (8) |
| GS1135 | XL1-Blue, pUT18C-*pbp1*, $amp^r$ | (8) |
| GS1136 | XL1-Blue, pKT25-*pbp2*, $kan^r$ | (8) |
| GS1137 | XL1-Blue, pUT18C-*pbp2*, $amp^r$ | (8) |
| GS1138 | XL1-Blue, pKT25-*pbp3*, $kan^r$ | (8) |
| GS1139 | XL1-Blue, pUT18C-*pbp3*, $amp^r$ | (8) |
| GS1187 | XL1-Blue, pKNT25-*ezrA*, $kan^r$ | (8) |
| GS1188 | XL1-Blue, pUT18-*ezrA*, $amp^r$ | (8) |

**Table S3. Primers used in this study.**

| Primer | Sequence (5'-3')[a] |
|---|---|
| **Primers used for construction of plasmids used in subcellular screening** | |
| im1_linker-FP_F_BamHI | ACTGGATCCCCGGATCTGGTGGAGAAGCTGCA |
| im2_m(sf)gfp_R_NotI_ EcoRI | AGTGAATTCGCGGCCGCTTACTTATAAAGCTCATCCATGCC |
| im77_SA1908_F_SalI_RBS | ATCGTCGAC*CAATAAAACTAGGAGGAAATTTAA*ATGGATTTATCTTCACCGATAG |
| im78_SA1908_R_BamHI | TCCGGGGATCCAATTGAATGATTCAATTTTATCCATC |
| **Primers used for making pLOW-*dcas9* compatible in MRSA** | |
| im183_pLOW_F | TACTGCAATCGGATGCGATTA |
| im184_pLOW_R | GTTAAGGGATGCATAAACTGC |
| im185_aad9_F_ol-im183 | *TAATCGCATCCGATTGCAGTA*ATTGGGCCCACCTAGGATC |
| im186_aad9_R_ol-im184 | *GCAGTTTATGCATCCCTTAAC*GCCGCGGTAATAAACTATCAA |
| **Primers used for construction of sgRNA plasmids used in depletion strains** | |
| mk299_sgRNA_1908 | TACCTAAGGCAACTAAAAAA GTTTAAGAGCTATGCTGGAAACAG |
| mk323_sgRNA_01908_V2 | ATAATGAGTCCAATGACTAT GTTTAAGAGCTATGCTGGAAACAG |
| **Primers used for chromosomal fusions and plasmids for localization studies** | |
| im147_SA1908_up_F_MluI | ACCTACGCGTGATTTTCGGTATATAAATGATAA |
| im148_SA1908_R_NotI_flag-overlap | *TCTTTATAATCAATATCATGATCTTTATAATCACCATCATGATCTTTATAATCC*GCGGCCGCATTGAATGATTCAATTTTATCCA |
| im149_aad9_up_F_SpeI_flag-overlap | *GATCATGATATTGATTATAAAGATGATGATGATAAATAA*ACTAGTATTGGGCCCACCTAGGAT |
| im150_aad9_down_R_1908 down-overlap | *TCATCACTTCAGCCTAACAT*CTCGAGGCCGCGGTAAT |
| im151_SA1908_down_F | ATGTTAGGCTGAAGTGATGA |
| im152_SA1908_down_R_BamHI | AGTCGGATCCTGATTTAAACCATCAATTCGC |
| im153_linker-FP_F_NotI | ACTGCGGCCGCCGGATCTGGTGGAGAAGCTG |
| im154_m(sf)gfp_R_SpeI | AGTACTAGTTTACTTATAAAGCTCATCCATG |
| im5_mKate_R_NotI_EcoRI | AGTGAATTCGCGGCCGCTTAACGGTGTCCCAATTTACTAGG |
| USHC109 | GCGACGCGTTTAACGGTGTCCCAATTTACTAGG |
| USHC148 | CGCGTCGACAGGAGGATAATTATTTATGTTAGAATTTGAACAAGG |
| im3_cfp_myfp_R_NotI_EcoRI | AGTGAATTCGCGGCCGCTTATTTATAAAGTTCGTCCATACC |
| **Primers used for verifying *smdA* silencing** | |
| im126_RT-q_pta_F | ATCATTGATGGCGAATTCCAAT |
| im127_RT-q_pta_R | GGACCAACTGCATCATATCC |
| im137_RT-q_SA1908_F | TATGTAACGGACATGAGATTAAT |
| im138_RT-q_SA1908_R | CTAATACCATTATAAACATGACC |

| | |
|---|---|
| **Primers used for construction of plasmids for overexpression** | |
| mk517_1908_R_NotI | ACGAGCGGCCGCCATATAGTCATCACTTCAGCCT |
| mk518_1908_F_RBS_SalI | ATCCAGTCGAC*CAATAAAACTAGGAGGAAATTTAA*ATG |
| | AGTAAGAAAAAAGTTAAGCGACAAAC |
| mk519_1908_H145A_F | GAACGAATTAGT**GC**TTTAGTATTAACAAG |
| mk520_1908_H145A_R | CTTGTTAATACTAAA**GC**ACTAATTCGTTC |
| mk521_1908_RT_AA_F | TTAGTATTAACA**GC**A**G**CTGGTCTTTATATT |
| mk522_1908_RT_AA_R | AATATAAAGACCAG**CTGC**TGTTAATACTAA |
| mk529_1908_FH_AA_F | CTTTAACAAATTTGTA**GC**C**GC**TGGTCGTATTCAAT |
| mk530_1908_FH_AA_R | ATTGAATACGACCA**GC**G**GC**TACAAATTTGTTAAAG |
| **Primers used for construction of plasmids to BACTH assays** | |
| gs718_SA1908_F_BamHI | GATCGGATCCCGATTTATCTTCACCGATAGTCATTG |
| gs719_SA1908_R_KpnI | GATCGGTACCCGATTGAATGATTCAATTTTATCCATC |
| gs735_SA1908ΔTMH_F_BamHI | GATCGGATCCCGGTAGTAAGAAAAAAGTTAAGCGAC |

[a]Restriction sites are underlined, sequences included as overhang in italic and inserted mutations in bold.
